# *Cask* dysfunction in inhibitory neurons reveals distinct synaptic and epileptic phenotypes in mouse models of *CASK*-related disorder

**DOI:** 10.64898/2026.11.17.767354

**Authors:** Kevin Jiang, Dongwon Lee, Lu-Tang Yang, Zhuohui Luo, Sty Karatayev, Nikhita Menon, Bhargav Vaidya, Sophie Medico, Mingshan Xue

**Affiliations:** Department of Neuroscience, Baylor College of Medicine, Houston, Texas 77030, USA; The Cain Foundation Laboratories, Jan and Dan Duncan Neurological Research Institute at Texas Children’s Hospital, Houston, Texas 77030, USA; Department of Molecular and Human Genetics, Baylor College of Medicine, Houston, Texas 77030, USA

## Abstract

Epilepsy is a major neurological manifestation of genetic neurodevelopmental disorders, yet how genetic disruptions produce distinct seizure phenotypes remains poorly understood. *CASK*-related disorder is an X-linked neurodevelopmental disorder frequently accompanied by epilepsy. It is caused by pathogenic variants in *CASK*, which encodes calcium/calmodulin-dependent serine protein kinase, a protein with diverse molecular functions. Here, we establish sex-specific mouse models of *CASK*-related disorder and identify epileptic phenotypes in both sexes. Cell type-specific *Cask* deletion identifies inhibitory neurons as a critical cellular locus of distinct forms of epileptic activity. *Cask* deficiency in parvalbumin- or somatostatin-expressing inhibitory neurons primarily causes spike-wave discharges or motor seizures, accompanied by reduced spontaneous or evoked inhibitory synaptic transmission, respectively. Thus, *Cask* dysfunction in different inhibitory synapses and circuits produces distinct seizure phenotypes. These findings reveal how cell type-specific synaptic dysfunction shapes epilepsy in *CASK*-related disorder and may provide insights into diverse neurological phenotypes of neurodevelopmental disorders.

## INTRODUCTION

Genetic neurodevelopmental disorders (NDDs) are a major cause of epilepsy (Oliver et al., 2023; Delgado-Escueta, 2024; Scorrano et al., 2024). For many genes implicated in these disorders, the molecular interactions and cellular functions of their encoded proteins have been extensively characterized. However, the pathological mechanisms underlying these disorders often remain unclear, as it remains challenging to bridge the gap from genetic deficit and cellular dysfunction to epilepsy and other neurological manifestations. This disconnect limits our understanding of the diseases and ability to develop mechanism-based therapeutics.

X-linked *CASK*-related disorder is a particularly informative example highlighting this knowledge gap. This severe NDD is caused by pathogenic variants in *CASK* and characterized by a spectrum of neurologic symptoms including global developmental delay, intellectual disability, microcephaly, pontocerebellar hypoplasia, neuromuscular and movement disorders, and epilepsy (Najm et al., 2008; Moog et al., 2011; Giacomini et al., 2021; Martin et al., 2025; Jiang et al., 2026a). The affected female heterozygous individuals predominantly carry genetic null variants, whereas male hemizygous individuals often have missense, partial loss-of-function variants, or somatic mosaicisms, because total loss of *CASK* is likely to cause early lethality (Moog et al., 2015; Jiang et al., 2026a). *CASK* encodes calcium/calmodulin-dependent serine protein kinase, which is expressed ubiquitously in the brain and has extensive molecular interactions across multiple cellular compartments. As a member of the membrane-associated guanylate kinase (MAGUK) family (Funke et al., 2005), CASK interacts through its multiple binding domains with numerous presynaptic proteins (Mori et al., 2023) including Neurexins, synaptic adhesion and specification molecules (Hata et al., 1996; Gomez et al., 2021), and Liprin-α proteins, key organizers of presynaptic sites (Wei et al., 2011; Spangler et al., 2013; Marcó de la Cruz et al., 2024). CASK also has postsynaptic and potentially nuclear functions such as scaffolding adhesion molecules (Hsueh et al., 1998; Chao et al., 2008; Gao et al., 2018), trafficking of inward rectifier potassium channels and NMDA receptors (Leonoudakis et al., 2004; Jeyifous et al., 2009), and regulation of gene expression (Hsueh et al., 2000; Wang et al., 2004; Mori et al., 2019). Consistent with these diverse molecular functions, loss of Cask/CASK in mouse or human neurons results in reduced inhibitory presynaptic markers (Becker et al., 2020), altered synaptic transmission (Atasoy et al., 2007; McSweeney et al., 2022), reduced NMDA receptor Grin2b levels in cortical pyramidal neurons (Mori et al., 2019), altered AMPA receptor GluA1 levels and localization in inhibitory neurons (Gao et al., 2019), and reduced dendritic arborization in inhibitory neurons (Gao et al., 2018). However, how these diverse cellular consequences of CASK dysfunction contribute to the distinct neurologic phenotypes of *CASK*-related disorder remain unclear.

Mouse models of *CASK*-related disorder are a critical system to begin linking these molecular and cellular functions to disease mechanisms *in vivo*. *CASK*-related disorder was modeled in mice by a null allele in heterozygous females and a hypomorph *tm1Sud*_allele in hemizygous males (Atasoy et al., 2007). Both models phenocopy microcephaly and pontocerebellar hypoplasia, in part due to the loss of cerebellar neurons (Najm et al., 2008; Srivastava et al., 2016; Patel et al., 2022; Guo et al., 2023; Mori et al., 2025). Female heterozygous knockout mice also show motor deficits (Srivastava et al., 2016). Other neurologic phenotypes, however, have not been comprehensively modeled or characterized. Epilepsy is of particular interest because it is one of the cardinal symptoms of *CASK*-related disorder, encompassing diverse seizure types (Takanashi et al., 2012; Giacomini et al., 2021; Jiang et al., 2026a), and its circuit-level mechanisms can be investigated in mouse models. Establishing and mechanistically dissecting epilepsy in mouse models of *CASK*-related disorder provides an opportunity to narrow the knowledge gap between genetic mechanism and neural circuit dysfunction and determine how *CASK* dysfunction produces a specific neurological phenotype.

To this end, we first characterized the epileptic phenotypes of sex-specific mouse models of *CASK*-related disorder and then used a novel *Cask* flox allele to selectively delete *Cask* in different neuronal types to determine the mechanisms underlying distinct seizure phenotypes.

## RESULTS

### Sex-specific mouse models of *CASK*-related disorder

To model null variants in *CASK*-related disorder, we generated a novel *Cask* flox allele (*tm1c*) with two *loxP* sites flanking exon 6 and crossed female heterozygous *Cask^tm1c/+^* mice with male hemizygous *Sox2-Cre^Tg/+^* mice, which express Cre in all epiblast cells (Hayashi et al., 2002) (Figure 1A,B). Consistent with the previous *Cask* knockout (KO) mice (Atasoy et al., 2007; Mori et al., 2019), male offspring *Sox2-Cre^Tg^*^/+^;*Cask^tm1c/y^*(Sox2-KO) mice died within the first day of life. Reverse transcription quantitative PCR (RT-qPCR) analysis targeting exons 8–11 of *Cask* showed that in the postnatal day 0 (P0) brain tissues, *Cask* mRNA levels in the male wildtype (WT), *Cask^tm1c/y^*, and *Sox2-Cre^Tg^*^/+^ mice were comparable, but reduced by 70% in Sox2-KO mice, indicating that the premature stop codon in exon 7 induced by the deletion of exon 6 (Figure 1A) leads to nonsense-mediated mRNA decay (Figure 1C). The residual mRNA in theory could produce a truncated Cask protein of 19.7 kDa, so we used two different antibodies to detect Cask by Western blots: one recognizing amino acid residues 318–415, the LIN2/LIN7 (L27) domain, after protein truncation and the other recognizing the N-terminal amino acids prior to protein truncation. Full-length Cask protein was similar among WT, *Cask^tm1c/y^*, and *Sox2-Cre^Tg^*^/+^ mice, but absent in Sox2-KO mice, validating that *Sox2-Cre* induced a complete recombination of the *tm1c* allele, resulting in a *Cask* KO *tm1d* allele (Figure 1A,D,E). Furthermore, Western blots using the N-terminal antibody did not detect a truncated Cask protein, indicating that any protein translated from the residual mRNA was degraded (Figure 1D).

**Figure 1:**
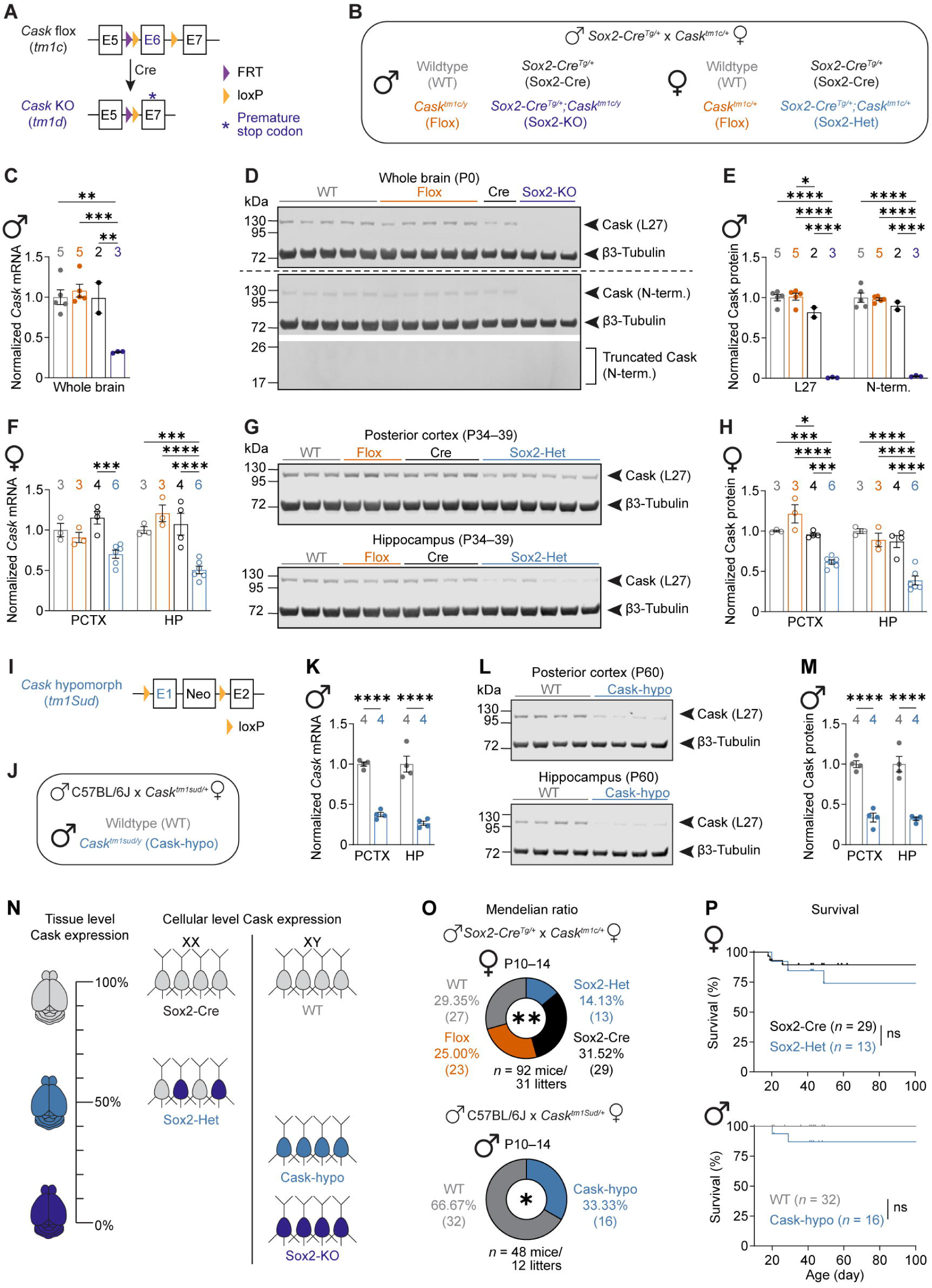
*Cask* heterozygous null and hemizygous hypomorphic alleles establish sex-specific mouse models of *CASK*-related disorder. **(A)** Genomic structures of *Cask* flox (*tm1c*) and KO (*tm1d*) allele. In the *tm1c* allele, Cre-mediated recombination between the two loxP sites deletes exon 6, resulting in the *tm1d* allele with a premature stop codon in exon 7. E, exon; FRT, Flp recombination site; loxP, Cre recombination site. **(B)** Offspring from *Sox2-Cre^Tg/+^* crossed with *Cask^tm1c/+^*. The color scheme is maintained across all panels for male and female mouse models separately. **(C)** Summary data of RT-qPCR analysis of transcripts from whole brain lysates of P0 males. *Cask* mRNA levels were reduced by 70% in *Sox2-Cre^Tg^*^/+^;*Cask^tm1c/y^*(Sox2-KO) mice as compared to wildtype (WT), *Cask^tm1c/y^* (Flox), and *Sox2-Cre^Tg^*^/+^ (Cre) mice. Data are mean ± S.E.M., one-way ANOVA with Tukey multiple comparison test. **(D)** Western blots of proteins from whole brain lysates of P0 males. Cask was detected via an antibody targeting the Cask L27 domain (top) or N-terminus (bottom). β3-Tubulin as loading control. The N-terminus antibody did not detect a truncated Cask translated from the residual *Cask* mRNA. **(E)** Summary data of normalized Cask protein levels from whole brain lysates of P0 males. Cask levels were first normalized by the β3-Tubulin levels and then by the average Cask levels of all WT mice from the same blots. Cask was absent in Sox2-KO mice. Data are mean ± S.E.M., two-way ANOVA with Tukey multiple comparison test. **(F–H)** Same as (C–E) but for posterior cortex (PCTX) and hippocampus (HP) in P34–39 females. *Cask* mRNA and Cask protein levels in *Sox2-Cre^Tg^*^/+^;*Cask^tm1c/+^*(Sox2-Het) mice were lower than those in WT, *Cask^tm1c/+^* (Flox), and *Sox2-Cre^Tg^*^/+^ (Cre) mice. Data are mean ± S.E.M., two-way ANOVA with Tukey multiple comparison test. **(I)** Genomic structure of *Cask* hypomorphic allele (*tm1Sud*), where exon 1 and a neomycin (Neo) cassette inserted in intron 1 are flanked by two loxP sites. **(J)** Male offspring from C57/BL6 crossed with *Cask^tm1Sud^*^/+^. The color scheme is maintained across all panels. **(K–M)** Same as (C–E) but for PCTX and HP in P60 males. *Cask* mRNA and Cask protein levels in Cask-hypo mice were reduced by 60–70% as compared to WT. Data are mean ± S.E.M., two-way ANOVA with Šídák’s multiple comparisons test. **(N)** Diagram illustrating the Cask expression at the tissue and cellular levels in different mouse models of *CASK*-related disorder. **(O)** Pie charts representing the observed numbers of mice with different genotypes at P10–14. Fewer Sox2-Het (top) and Cask-hypo mice (bottom) were observed than the Mendelian expectation. Chi-square test (Sox2-Het) or binomial test (Cask-hypo). **(P)** Neither Sox2-Het nor Cask-hypo mice showed survival differences after genotyping. Tic marks indicate censured mice that were euthanized for non-medical reasons. Since Sox2-Cre results in complete recombination of the *tm1c* to *tm1d* allele, only Sox2-Cre was used as control. Log-rank (Mantel-Cox) test. Filled (male) and open (female) symbols represent individual mice. The numbers (*n*) are indicated in the panels. ns, *P* > 0.05; *, *P* < 0.05; **, *P* < 0.01; ***, *P* < 0.001; ****, *P* < 0.0001.

At the time of genotyping (i.e., P10–14), female *Sox2-Cre^Tg^*^/+^;*Cask^tm1c/+^*(Sox2-Het) mice were 14% of the total female offspring, which is lower than the expected Mendelian frequency, indicating a partial early lethality phenotype (Figure 1O). However, no significant survival differences were observed following genotyping (Figure 1P). *Cask* mRNA and proteins levels in Sox2-Het mice were reduced by 30–50% and 40–60%, respectively, as compared to female WT, *Cask^tm1c/+^*, and *Sox2-Cre^Tg^*^/+^ mice at 1 month of age (Figure 1F–H). Thus, Sox2-Het mice are equivalent to constitutive heterozygous KO mice and serve as a female mouse model of *CASK*-related disorder.

To model the disorder in male mice we used the *Cask* hypomorphic *tm1Sud* allele, which contains two loxP sites flanking exon 1 and a neomycin resistance gene cassette in intron 1 (Atasoy et al., 2007) and has been reported to reduce Cask protein by 60–70% prior to Cre-mediated recombination (Atasoy et al., 2007; Srivastava et al., 2016). We crossed female heterozygous *Cask^tm1Sud/+^* mice with male WT C57BL/6J mice (Figure 1I,J). At the time of genotyping (i.e., P10–14), male *Cask^tm1Sud/y^* (Cask-hypo) mice were 33% of the total male offspring, indicating a partial early lethality phenotype (Figure 1O), but no significant survival differences were observed following genotyping (Figure 1P). *Cask* mRNA and protein levels in Cask-hypo mice were reduced by 60–70% and ∼70%, respectively, as compared to male WT littermates at 2 months of age, indicating that the disruptions to Cask expression occur at the transcriptional level (Figure 1K–M).

Across both models there is tissue level reduction of Cask; however, since *Cask* is an X-linked gene the two different models are unique in their cellular expression. In Sox2-Het mice, X-inactivation creates a mosaic condition where cells either express 100% Cask from the WT allele or 0% Cask from the KO allele, whereas in Cask-hypo mice all cells express ∼30% Cask from the hypomorphic allele (Figure 1N). Furthermore, we observed reduced body weight, microcephaly, and pontocerebellar hypoplasia in both Sox2-Het and Cask-hypo mice (Supplemental figure 1A–F), consistent with previous reports (Najm et al., 2008; Srivastava et al., 2016; Mori et al., 2019). Thus, Sox2-Het and Cask-hypo mice both molecularly show reductions of Cask expression and phenotypically exhibit gross anatomical and developmental features similar to individuals with *CASK*-related disorder. Hence, we next investigated epilepsy in both models.

### Sex-specific mouse models of *CASK*-related disorder exhibit spontaneous electrographic seizures

To record video-electroencephalography (EEG) and electromyography (EMG), we implanted intracranial electrodes over the left frontal cortex, bilateral somatosensory cortices, and an EMG electrode in the neck muscles at P35–42. Following recovery, mice underwent 2–4 recording sessions (48–72 hours per session) spaced 6–8 weeks apart. For Sox2-Het mice, only age-matched female *Sox2-Cre^Tg^*^/+^ (Sox2-Cre) littermates were used as the control to account for the *Sox2-Cre* transgene, as the Cask flox allele (*tm1c*) is completely converted to the KO allele (Figure 1D,E). For Cask-hypo mice, age-matched male WT littermates were used as the control.

Both Sox2-Het and Cask-hypo mice exhibited spontaneous spike-wave discharges (SWDs), a form of non-motor epileptic activity that parallels human absence seizures and is frequently observed in animal models of epilepsy (Maheshwari and Noebels, 2014). Behavioral arrest frequently co-occurred during these electrographic events as evidenced by the reduction of EMG activity (Figure 2A). The frequencies of SWDs in both Sox2-Het and Cask-hypo mice were significantly higher than those in control mice across all recording timepoints (Figure 2B). SWDs occurred throughout the day and night, and the frequency peaked shortly after dark onset in both models (Figure 2C). Ethosuximide, the first-line treatment for absence seizures in humans, suppressed these SWDs (Figure 2D), indicating the involvement of T-type calcium channels in these SWDs that is commonly observed in other models of absence epilepsy (Maheshwari and Noebels, 2014).

**Figure 2:**
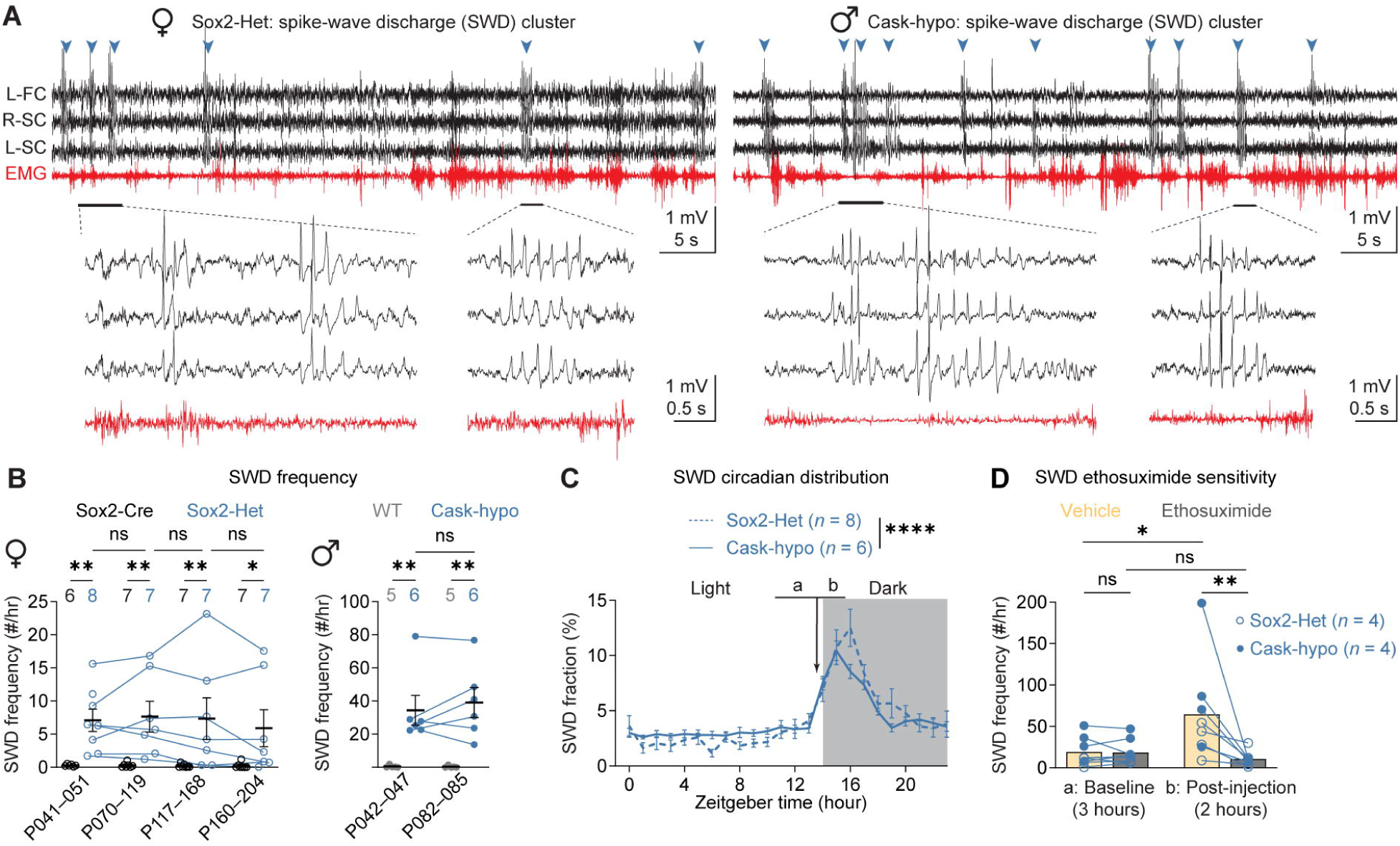
Sex-specific mouse models of CASK-related disorder show spontaneous spike-wave discharges (SWDs). **(A)** EEG/EMG traces from a Sox2-Het (left) or Cask-hypo (right) mouse experiencing a cluster of SWDs indicated by arrow heads. Four SWDs are expanded below highlighting the morphology and spike frequency of 6–8 Hz. L-FC, left frontal cortex; R-SC, right somatosensory cortex; L-SC, left somatosensory cortex; EMG, from neck muscles. **(B)** Summary data showing that the SWD frequencies of Sox2-Het (left) and Cask-hypo (right) were significantly increased as compared to their respective control mice at different recording ages, with no differences between each recording. Data are mean ± S.E.M., mixed-effect analysis with Fisher’s LSD test (Sox2-Het) or two-way ANOVA with Fisher’s LSD test (Cask-hypo). **(C)** The percent of SWDs in each hour over a day is plotted as a function of the zeitgeber time (ZT) and averaged across the four days of a recording and the first two recording timepoints if possible, to display the circadian distribution of SWDs in Sox2-Het and Cask-hypo mice. Gray background indicates the dark period (ZT14–24). 59.13% of SWD frequency variance was explained by ZT hour for Sox2-Het and Cask-hypo together (*P* < 0.0001). Data are mean ± S.E.M., two-way ANOVA. **(D)** Sox2-Cre and Cask-hypo mice were injected with ethosuximide (200 mg/kg body weight) and vehicle (PBS), each on one of the two consecutive recording days at the ZT hour 13:30 as indicated by the arrow in (C). SWD frequences were similar during the 3-hour baseline period prior to the injection as indicated by the ‘a’ in (C). During the 2-hour post-injection period as indicated by the ‘b’ in (C), SWD frequency was significantly lower for ethosuximide than vehicle. Columns indicate mean, two-way ANOVA. Filled (male) and open (female) symbol represents individual mice. The numbers (*n*) are indicated in the panels. ns, *P* > 0.05; *, *P* < 0.05; **, *P* < 0.01.

In addition to SWDs, we observed that Sox2-Het mice, but not Cask-hypo mice, on rare occasion exhibited clonus and wild running, although no such event was captured in the video-EEG/EMG recordings. Those mice that exhibited these events often died after the seizures. Thus, our studies establish that mouse models of *CASK*-related disorder have epilepsy, majorly in the form of spontaneous SWDs.

### *Cask* deletion in inhibitory but not excitatory neurons causes severe juvenile-onset epilepsy

To understand the cellular origins of epilepsy in *CASK*-related disorder, we used the flox *tm1c* allele and cell type-specific Cre lines to delete *Cask* in neuronal types of interest, while leaving *Cask* expression intact in all other cells. We focused these cell type-specific conditional KO (cKO) studies on male hemizygous instead of female heterozygous mice for three reasons: (1) in humans, males with *CASK*-related disorder have a higher penetrance of epilepsy that is more severe and earlier in onset— especially so amongst those with loss-of-function variants (Jiang et al., 2026a); (2) in mice, male Cask-hypo mice have a higher frequency of SWDs than female Sox2-Het mice (Figure 2B); and (3) the single X-chromosome in males allows *Cask* deletion to uniformly affect the population of interest, rather than result in a variable and mosaic penetrance. Importantly, video-EEG/EMG recordings showed that unlike the *Cask^tm1Sud/y^*mice, the new flox mice (*Cask^tm1c/y^*) did not show increased SWDs or other forms of epileptic activity as compared to WT mice (Supplemental figure 3A), validating their use for the cKO studies.

We initially targeted two broad categories of neurons in the brain, excitatory neurons by pan-glutamatergic *Vglut2-ires-Cre* and inhibitory neurons by pan-GABAergic/glycinergic *Viaat-ires-Cre* (Vong et al., 2011; Kim et al., 2024). We crossed female heterozygous *Cask^tm1c/+^*mice with male heterozygous *Vglut2^Cre/+^* or *Viaat^Cre/+^*mice (Supplemental figure 2A) and found near absence of male hemizygous *Vglut2^Cre/+^*;*Cask^tm1c/y^* (Vglut2-cKO) or *Viaat^Cre/+^*;*Cask^tm1c/y^* (Viaat-cKO) mice, respectively, on P10 (Supplemental figure 2B,C). In cases where the litters were monitored from birth, both Vglut2-cKO and Viaat-cKO mice were typically found dead on P0–1. Two Viaat-cKO pups survived until P14 and were observed to exhibit convulsive activity (Supplemental figure 2D). Thus, these results demonstrate a vital role of *Cask* in both neuronal populations to mouse survival.

To reduce lethality, we next used *Vglut1-ires-Cre* or *Emx1-ires-Cre* to delete *Cask* from the excitatory neurons in the cortex and hippocampus (Gorski et al., 2002; Harris et al., 2014) by crossing female heterozygous *Cask^tm1c/+^* mice with male homozygous *Vglut1^Cre/Cre^* or *Emx1^Cre/Cre^*mice (Figure 3A). Both *Vglut1^Cre/+^*;*Cask^tm1c/y^*(Vglut1-cKO) and *Emx1^Cre/+^*;*Cask^tm1c/y^*(Emx1-cKO) mice were observed at the expected Mendelian ratio at P10–14 and exhibited no post-genotyping survival differences (Figure 3B,C). Neither cKO mice exhibited increased SWDs or other forms of generalized epileptic activity (Figure 3D; Supplemental figure 3B).

**Figure 3:**
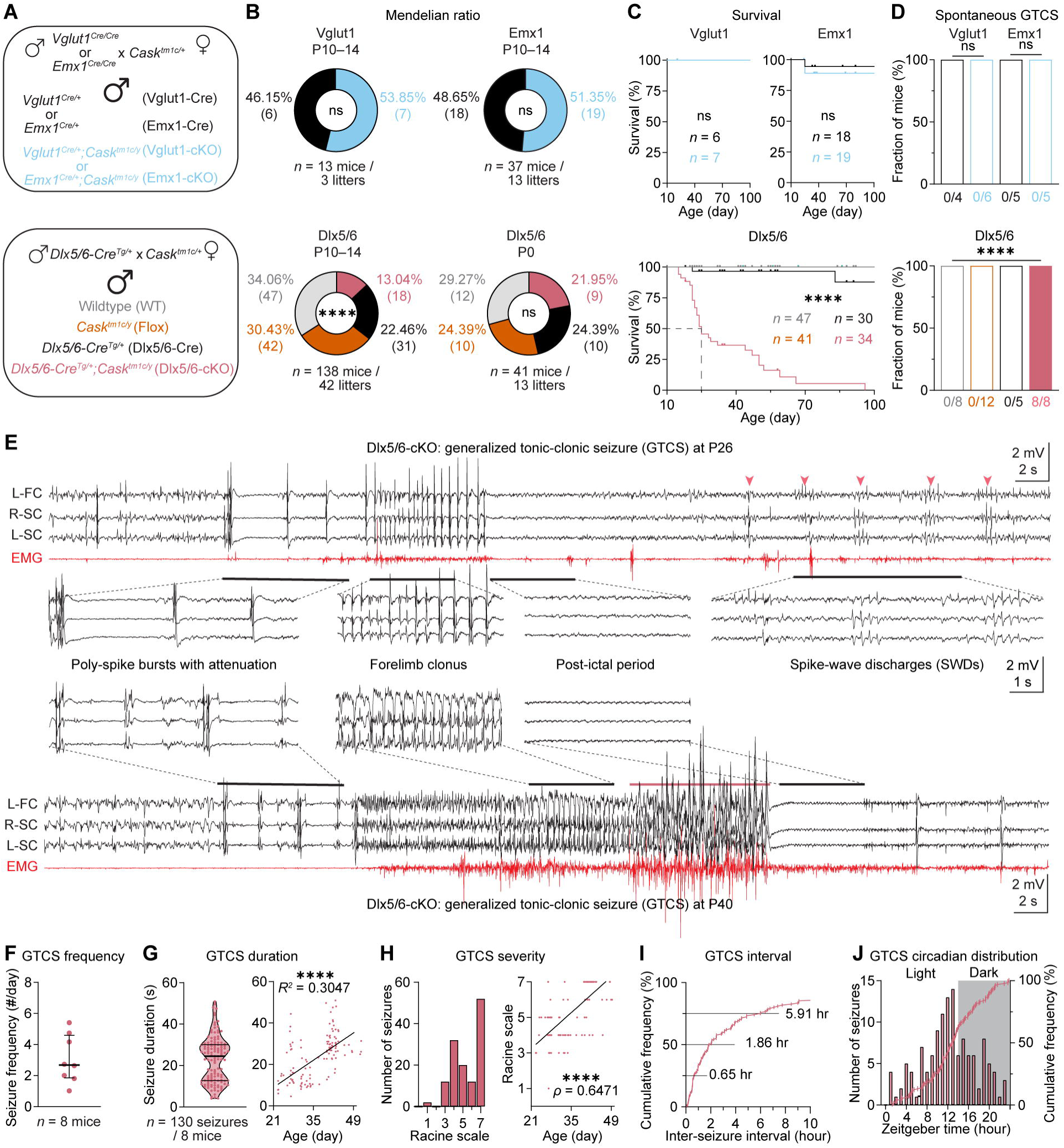
*Cask* deletion in forebrain inhibitory neurons results in premature lethality and juvenile epilepsy. **(A)** Male offspring from *Vglut1^Cre/Cre^, Emx1^Cre/Cre^*, or *Dlx5/6^Cre/+^* crossed with *Cask^tm1c/+^*. The color scheme is maintained across all panels. **(B)** Pie charts representing the observed numbers of mice with different genotypes at P10–14 or P0. Vglut1-cKO and Emx1-cKO mice showed expected Mendelian ratios, while fewer Dlx5/6-cKO mice were observed than the Mendelian expectation at P10–14, but not at P0. Binomial test (Vglut1-cKO and Emx1-cKO) or Chi-square test (Dlx5/6-cKO). **(C)** Neither Vglut1-cKO nor Emx1-cKO mice showed survival differences compared to controls. Dlx5/6-cKO mice exhibited marked early lethality with a median survival of P25. Tic marks indicate censured mice that were euthanized for non-medical reasons. Log-rank (Mantel-Cox) test with Holm-Šídák’s multiple comparisons test. **(D)** No Vglut1-cKO or Emx1-cKO mice had a spontaneous generalized tonic-clonic seizure (GTCS) in any video-EEG/EMG recording. Dlx5/6-cKO mice, but not their littermate controls, all had GTCSs. Fisher’s exact test. **(E)** EEG/EMG traces from a Dlx5/6-cKO mouse experiencing a GTCS at P26 or P40. The expanded traces highlight EEG poly-spike bursts with attenuation that preceded a generalized seizure, the epileptic oscillations during forelimb clonus events, and the post-ictal period. In the P40 trace, after a wild-running phase (red line) the post-ictal period was associated with an attenuated baseline, theta oscillations, and tonic extension, while at P26 there were normal behaviors, and EEG returned to baseline. The P26 trace also captured a cluster of SWDs indicated by arrow heads. **(F–J)** Summary data of GTCS features in Dlx5/6-cKO mice. **(F)** GTCS frequency of each mouse across all recordings. Lines indicate median (2.683 GTCSs per day) ± interquartile range (I.Q.R., 1.022–4.583). **(G)** Left, violin plot displaying the duration of every GTCS, which shows a bi-modal distribution. Lines indicate median (24.48 s) ± I.Q.R. (12.73–30). Right, scatter plot showing a positive correlation between seizure duration and mouse age (Pearson’s correlation, *R^2^* = 0.3047). **(H)** Left, histogram of GTCS severity measured with a modified Racine score. Right, scatter plot showing a positive correlation between Racine score and mouse age (Spearman’s rank correlation, ρ = 0.6471). **(I)** Cumulative distribution of the inter-seizure interval. 17/116 (14.66%) of intervals are greater than 10 hours and not displayed. **(J)** Histogram showing the numbers of GTCSs in the ZT hours (left y axis) is overlayed with the cumulative distribution (right y axis). A high percentage of GTCSs (53 out of 130 seizures, 40.76%) occurred in ZT hours 9–13. Each symbol (*n*) represents a mouse (F) or seizure (G–J). The numbers (*n*) are indicated in the panels. ns, *P* > 0.05; ****, *P* < 0.0001.

In contrast, when we used *Dlx5/6-Cre* (Monory et al., 2006) to delete *Cask* from the forebrain inhibitory neurons by crossing female heterozygous *Cask^tm1c/+^* mice with male hemizygous *Dlx5/6-Cre^Tg/+^*mice (Figure 3A), the number of *Dlx5/6-Cre^Tg/+^*;*Cask^tm1c/y^*(Dlx5/6-cKO) mice was significantly below the Mendelian expectation at P10–14. However, Dlx5/6-cKO mice were observed at the expected Mendelian ratio at P0 (Figure 3B), indicating that a fraction of Dlx5/6-cKO mice die prematurely within the first two weeks of life. The remaining Dlx5/6-cKO mice continued to show significantly decreased survival, with a median lifespan of 25 days (Figure 3C).

We recorded video-EEG/EMG from Dlx5/6-cKO mice, as early as P21 with repeat recordings on the interval of several days due to their early lethality, and observed that each of the 8 recorded mice, but no littermate controls, experienced generalized tonic-clonic seizures (GTCSs) (Figure 3D,E). Additionally, other marked EEG abnormalities including high frequency of poly-spikes, continuous slow spike-wave during nonrapid eye movement (NREM) sleep, SWDs, and periods of clustered myoclonus were observed (Figure 3E; Supplemental Figure 4B,D–F). Notably littermate controls did not exhibit any similar EEG abnormalities (Supplemental Figure 4A,C). For GTCSs, the most severe form of epileptic activity, Dlx5/6-cKO mice had 2.683 GTCS per day (1.022–4.583, median ± interquartile range (I.Q.R.)) (Figure 3F) with a median seizure duration of 24.48 seconds (12.73–30 seconds, median ± I.Q.R.) (Figure 3G). Seizure duration increased with age (linear regression, *R^2^* = 0.3047, *P* < 0.0001) (Figure 3G), as well as the severity that was classified on a modified Racine scale (Van Erum et al., 2019) (Spearman’s rank correlation, ρ = 0.6471, *P* < 0.0001) (Figure 3H). While there were typically only a few seizures recorded per day, the inter-seizure interval was 1.86 hours (0.65–5.91 hours, median ± I.Q.R.), suggesting that seizures happened within temporal clusters (Figure 3I). Furthermore, GTCS preferentially occurred during Zeitgeber time (ZT) hours 9–14 (53 out of 130 seizures, 40.76%) (Figure 3J). Together, these results show that selective loss of Cask in the forebrain inhibitory neurons, but not cortical excitatory neurons, of male mice is sufficient to cause epilepsy.

### *Cask* deletion in different inhibitory neurons leads to distinct epileptic phenotypes

Inhibitory neurons in the forebrain consist of diverse subtypes that regulate distinct circuit elements of inhibition (Kepecs and Fishell, 2014; Tremblay et al., 2016; Feldmeyer et al., 2018; Huang and Paul, 2019). To further determine how different inhibitory circuits contribute to epilepsy in *CASK*-related disorder, we selectively deleted *Cask* in each of the three largely nonoverlapping interneuron populations, parvalbumin (Pv), somatostatin (Sst), or serotonin receptor 3a (Htr3a)-expressing cells utilizing the well-established *Pv-ires-Cre*, *Sst-ires-Cre*, or *Htr3a-Cre* lines, respectively (Figure 4A) (Rudy et al., 2011; Tremblay et al., 2016). We recorded video-EEG/EMG beginning around P42 with follow up intervals as described above. *Pv^Cre/+^*;*Cask^tm1c/y^* (Pv-cKO) mice showed SWDs that were typically 6–8 Hz and correlated with behavioral arrests (Figure 4B). The SWD frequency increased significantly with age and was higher than age-matched littermate controls at all recording timepoints (Figure 4C). The circadian rhythm of SWD frequency was similar to those of Sox2-Het and Cask-hypo mice and did not change with age despite the vast increase in SWD frequency (Figure 4D). These SWDs were also suppressed by ETX (Figure 4E). Additionally, Pv-cKO mice exhibited pronounced changes of NREM sleep with age (Supplemental figure 5), and two Pv-cKO mice exhibited a total 4 spontaneous GTCSs at the oldest recording timepoint (Figure 4F,G).

**Figure 4:**
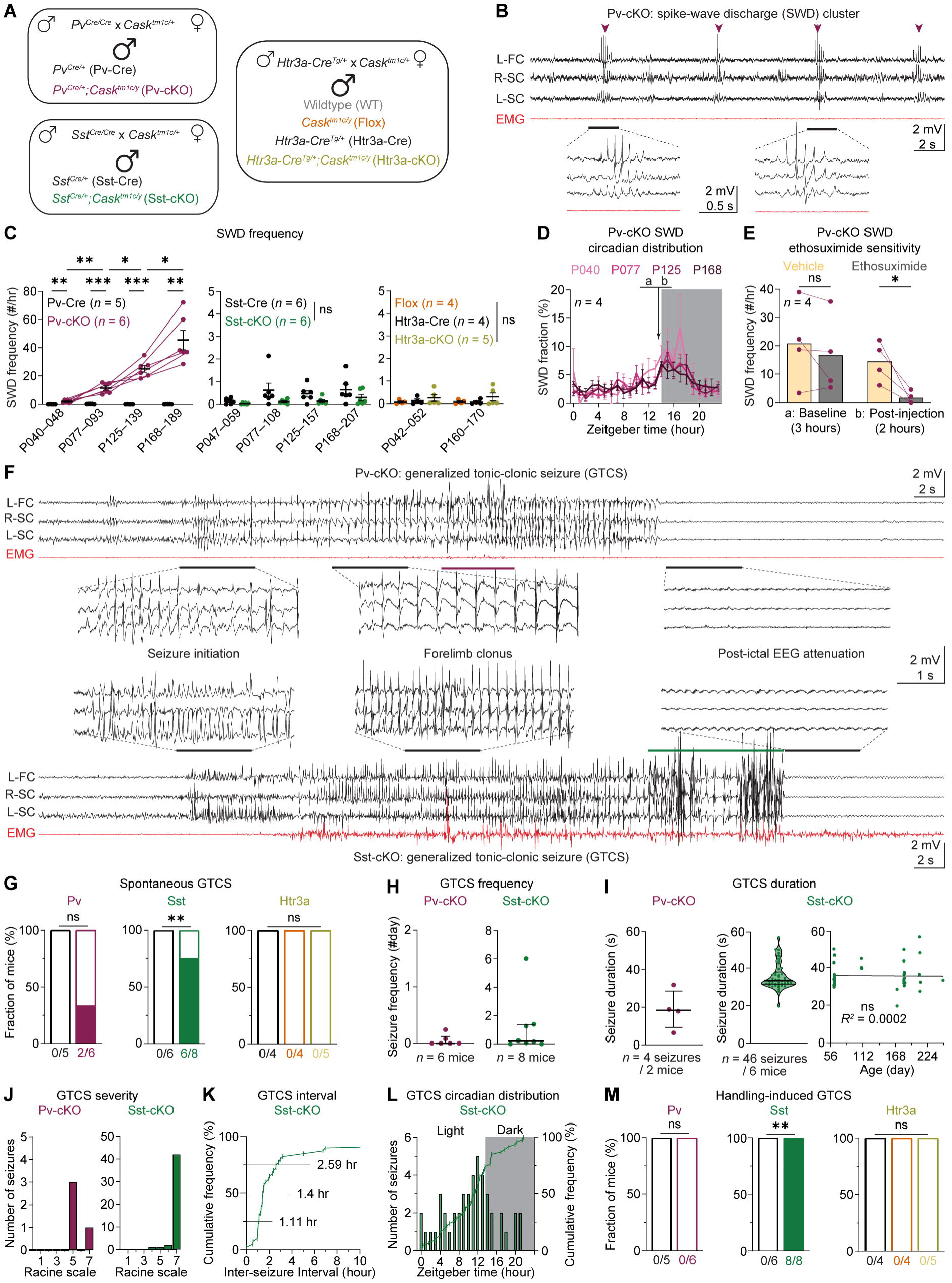
*Cask* deletion in Pv or Sst interneurons results in distinct epileptic phenotypes. **(A)** Male offspring from *Pv^Cre/Cre^*, *Sst^Cre/Cre^*, or *Htr3a-Cre^Tg/+^* crossed to *Cask^tm1c/+^*. The color scheme is maintained across all panels. **(B)** EEG/EMG traces from a Pv-cKO mouse showing a cluster of SWDs indicated by arrow heads. Two SWDs are expanded below highlighting the SWD morphology and spike frequency of 6–8 Hz. **(C)** Summary data of SWD frequency for Pv-cKO, Sst-cKO, Htr3a-cKO, and their respective control mice at different recording ages. Pv-cKO mice exhibited significantly increased frequencies at all timepoints, with significant increases between each recording. Data are mean ± S.E.M., two-way ANOVA with Tukey multiple comparison test. **(D)** The percent of SWDs in each hour over a day is plotted as a function of the ZT hour and averaged over two days of a recording to display the circadian distribution of SWDs in Pv-cKO mice. Each line represents the average distribution of 6 mice at each recording timepoint. Gray background indicates the dark period (ZT hours 14–24). 26.51% of SWD frequency variance was explained by ZT hour for four recording timepoints (*P* = 0.0078). Despite the increase of SWD frequency with age, the circadian distribution remains similar across four recording timepoints (*P* > 0.05). Data are mean ± S.E.M., two-way ANOVA. **(E)** Pv-cKO mice were injected with ethosuximide (200 mg/kg body weight) and vehicle (PBS), each on one of the two consecutive recording days at the ZT hour 13:30 as indicated by the arrow in (D). SWD frequences were similar during the 3-hour baseline period prior to the injection as indicated by the ‘a’ in (D). During the 2-hour post-injection period as indicated by the ‘b’ in (D), SWD frequency was significantly lower for ethosuximide than vehicle. Columns indicate mean, two-way ANOVA. **(F)** EEG/EMG traces from a Pv-cKO mouse (top) and an Sst-cKO mouse (bottom) experiencing a GTCS. The expanded traces highlight the differences in the EEG oscillations at the seizure initiation, during the forelimb clonus, and the immediate post-ictal state during tonic-extension. Purple (Pv) and green (Sst) lines highlight the period of wild-running. **(G)** An increased proportion of Sst-cKO, but not Pv-cKO and Htr3a-cKO, mice had spontaneous GTCSs during video-EEG/EMG recordings compared to controls. **(H–L)** Summary data of GTCS features in Pv-cKO and Sst-cKO mice. **(H)** GTCS frequency of each mouse across all recordings. Lines indicate median (Pv-cKO, 0.0 GTCSs per day; Sst-cKO, 0.200 GTCSs per day) ± I.Q.R. (Pv-cKO, 0.0–0.122; Sst-cKO, 0.019–1.360). **(I)** Left and middle, durations of individual GTCSs. Lines indicate median (Pv-cKO, 18.36 s; Sst-cKO, 33.21 s) ± I.Q.R. (Pv-cKO, 9.28–28.56; Sst-cKO, 31.39–37.87). Right, scatter plot showing no correlation between seizure duration and Sst-cKO mouse age (Pearson’s correlation, *R^2^* = 0.0002). **(J)** Histogram of GTCS severity measured with a modified Racine score. Most GTCSs in Pv-cKO mice were scored a 5 (clonus while lying down), while nearly all GTCSs in Sst-cKO mice reached a score of 7 (wild-running with tonic-extension). **(K)** Cumulative distribution of the inter-seizure interval for Sst-cKO mice. 4/40 (10%) of intervals are greater than 10 hours and not displayed. **(L)** Histogram showing the number of GTCSs in the ZT hours (left y axis) is overlayed with the cumulative distribution (right y axis) for Sst-cKO mice. A high percentage of GTCSs (19/46 seizures, 41.30%) occurred in ZT hours 9–13. **(M)** Only Sst-cKO mice exhibited handling-induced seizures. Each symbol represents a mouse (C,E,H) mice or seizure (I–L). The numbers (*n*) are indicated in the panels. ns, *P* > 0.05; *, *P* < 0.05; **, *P* < 0.01; ***, *P* < 0.001.

In contrast, neither *Sst^Cre/+^*;*Cask^tm1c/y^*(Sst-cKO) nor *Htr3a-Cre^Tg/+^*;*Cask^tm1c/y^* (Htr3a-cKO) mice had elevated SWDs (Figure 4C). However, spontaneous GTCSs were captured in one of the multiple video-EEG/EMG recordings for 6 of 8 recorded Sst-cKO mice (Figure 4F,G), with the earliest GTCSs observed at P60. Across all recordings, Sst-cKO mice had 0.1998 GTCS per day (0.01943–1.360, median ± I.Q.R.) (Figure 4H). The duration of GTCSs had a unimodal distribution with a median of 33.21 seconds (31.39–37.87 seconds, median ± I.Q.R.) and did not correlate with age (Figure 4I). The vast majority of the GTCSs (42/46, 91.3%) resulted in a modified Racine grade of 7 (wild running with tonic extension) (Figure 4J). Similar to Dlx5/6-cKO mice, Sst-cKO mice exhibited seizure clusters with an inter-seizure interval of 1.43 hours (1.11–2.76 hours, median ± I.Q.R.) (Figure 4K) and showed a higher frequency during ZT hours 9– 14 (22/46 seizures, 47.83%) (Figure 4L). While clinically displaying clonus, wild running, and post-ictal attenuation, the EEG patterns at the GTCS initiation and progression were markedly different between Sst-cKO and Pv-cKO mice (Figure 4F), suggesting differences in epileptic circuit dynamics. Video-EEG/EMG recordings from five Htr3a-cKO mice did not capture any GTCSs (Figure 4G). We also did not observe GTCSs in any littermate controls (Figure 4G). Finally, all Sst-cKO mice, but none of Pv-cKO and Htr3a-cKO mice, exhibited handling-induced GTCSs that began around 8–9 weeks of age (Figure 4M).

Altogether, these studies reveal that loss of Cask in different inhibitory neuron populations results in distinct epileptic phenotypes. To further understand the cellular mechanism of inhibitory dysfunction that contributes to these different seizures, we next examined the synaptic transmission, neuronal excitability, and densities of Pv and Sst interneurons in the primary somatosensory cortex (S1) where we recorded EEG.

### *Cask* deletion in Pv interneurons reduces spontaneous GABAergic transmission onto pyramidal neurons

At the earliest EEG recording timepoint, Pv-cKO mice exhibited low levels of SWDs (Figure 4C). Thus, we prepared acute brain slices at P42–56 and conducted electrophysiology experiments in artificial cerebrospinal fluid (ACSF) to identify changes that are most likely the primary alterations due to loss of Cask rather than potential circuit re-organization secondary to seizures. All mice carried a Cre-dependent EYFP reporter, *Rosa26-CAG-LSL-EYFP* (Ai3), for visualizing Pv interneurons. Given the potential presynaptic and postsynaptic roles of Cask, we performed whole-cell patch clamp recordings from pairs of a Pv interneuron and an adjacent pyramidal neuron that were within 50 μm to each other in layer 2/3 of the primary somatosensory cortex (S1) (Figure 5A). Presynaptic Pv interneurons were stimulated to fire a train of six action potentials at 10 Hz, and the resulting GABA release was measured by unitary inhibitory postsynaptic currents (uIPSCs) from the pyramidal neurons (Figure 5B). The connectivity rate, connection strength measured by the first uIPSC amplitude, or short-term synaptic depression did not show significant differences between control (*Pv^Cre/+^;Rosa26^Ai3/+^*) and Pv-cKO (*Pv^Cre/+^;Cask^tm1c/y^;Rosa26^Ai3/+^*) mice (Figure 5C–E).

**Figure 5:**
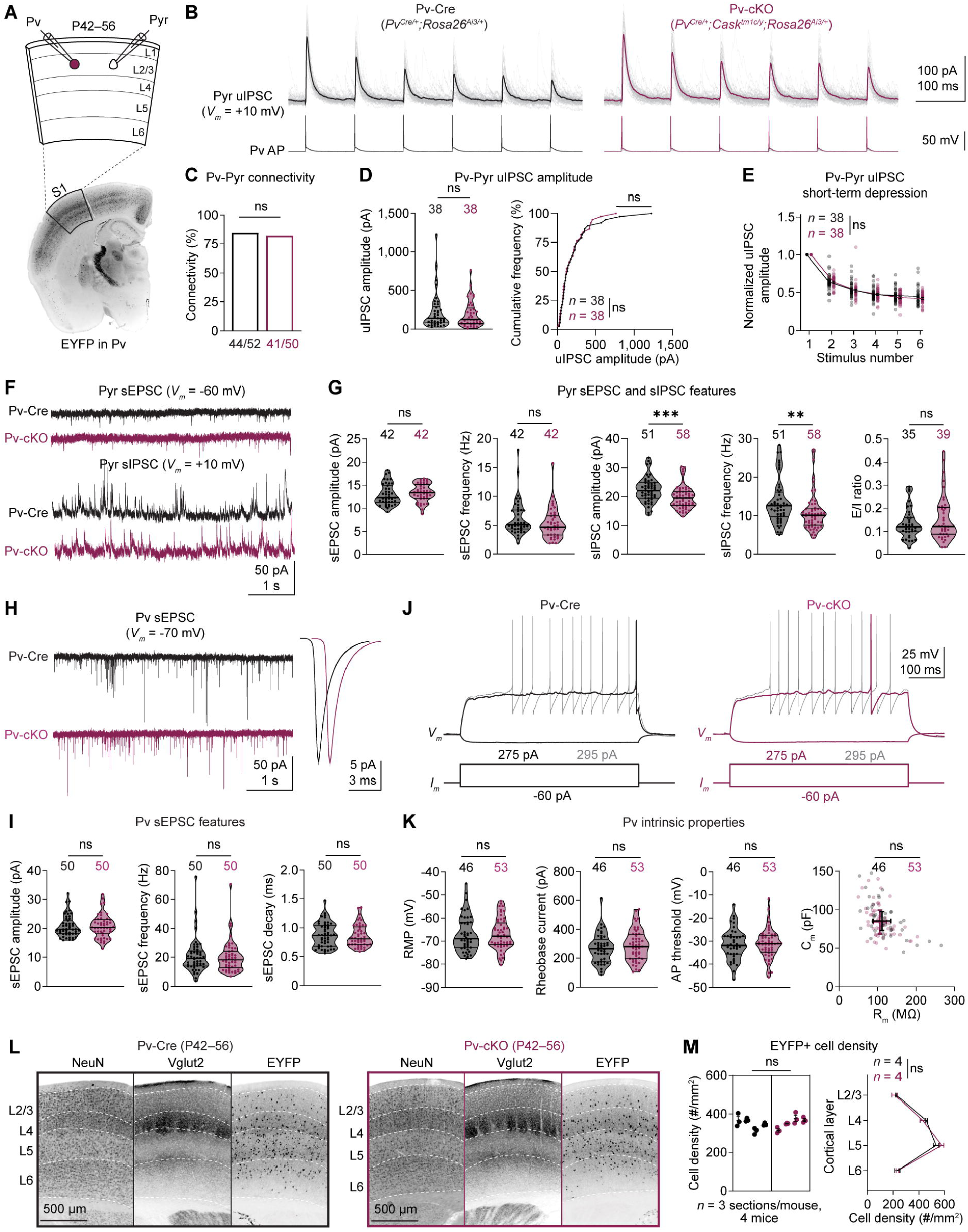
*Cask* deletion in Pv interneurons reduces global inhibition onto layer 2/3 pyramidal neurons. **(A)** Schematic of slice experiments (top), where a pair of adjacent (< 50 μm) Pv interneuron and pyramidal neuron were recorded in the layer 2/3 of the primary somatosensory cortex (S1) from 12 control Pv-Cre and 14 Pv-cKO male mice at P42– 56. Pv interneurons were identified by the Cre-dependent expression of EYFP (bottom). L, layer; Pyr, pyramidal neuron. **(B)** A train of six action potentials (APs) at 10 Hz in a Pv interneuron (lower panels) evoked uIPSCs in a pyramidal neuron held at +10 mV (upper panels). 30–50 individual traces (gray) and the average trace (color) are superimposed. **(C)** Unitary connection rates, Fisher’s exact test. Note, some connected pairs were excluded from the uIPSC analysis in (D,E) due to series resistances > 20 MΩ. **(D)** Left, summary data of uIPSC amplitudes evoked by the first action potentials in the train. Lines indicate median (Pv-Cre, 133.8 pA; Pv-cKO, 119.3 pA) ± I.Q.R., Mann-Whitney rank test. Right, cumulative frequencies of uIPSC amplitude. Kolmogorov-Smirnov test. **(E)** uIPSC amplitudes during the train of action potentials were normalized by the amplitudes of the first uIPSCs. Data are mean ± S.E.M., two-way ANOVA. **(F)** Representative traces of sEPSCs (top) and sIPSCs (bottom) from pyramidal neurons. **(G)** Summary data of the sEPSC amplitude, frequency, sIPSC amplitude, frequency, and E/I ratio of pyramidal neurons. Both the sIPSC amplitude and frequency were reduced in Pv-cKO mice. Lines indicate median ± I.Q.R., Mann-Whitney rank test. **(H)** Left, representative traces of sEPSCs from Pv interneurons. Right, the average sEPSC waveforms across all neurons from Pv-Cre (black) or Pv-cKO (purple) mice. **(I)** Summary data of sEPSC amplitude, frequency, and single-exponential decay time constant of Pv interneurons. Lines indicate median ± I.Q.R., Mann-Whitney rank test. **(J)** Membrane potentials (upper panels) in response to a hyperpolarizing (color), the rheobase (color), and a supra-rheobase (gray) current injections in Pv neurons. **(K)** Summary data of resting membrane potential (RMP), rheobase current, action potential threshold, membrane capacitance (C_m_), and membrane resistance (R_m_) of Pv interneurons. Lines indicate median ± I.Q.R., Mann-Whitney rank test. **(L)** Representative fluorescence images of brain sections stained for NeuN, Vglut2, and EYFP from Pv-Cre and Pv-cKO mice. **(M)** Left, EYFP+ cell density of a 1000-μm-wide cortical column in S1 barrel cortex. Each symbol represents a section, and each column represents a mouse. Lines indicate mean (of sections) ± S.E.M., nested t-test. Right, EYFP+ cell density in each cortical layer, calculated as an average of the 3 sections. Data are mean (of mice) ± S.E.M., two-way ANOVA. Each symbol represents a synaptic connection or neuron, unless otherwise specified. The numbers (*n*) are indicated in the panels. ns, *P* > 0.05; **, *P* < 0.01, ***, *P* < 0.001.

During these experiments, we also recorded spontaneous excitatory and inhibitory postsynaptic currents (sEPSCs and sIPSCs), a measure of global synaptic inputs onto a cell, from the pyramidal neurons as well as sEPSCs from the Pv interneurons. The median peak amplitude and frequency of sIPSCs, but not sEPSCs, of pyramidal neurons were modestly reduced by 11.7% and 19.9%, respectively, in Pv-cKO mice, revealing a reduction of inhibition. However, the overall excitatory/inhibitory (E/I) ratio, which was measured by the ratio between sEPSC and sIPSC charges per unit time, was not significantly altered (Figure 5F,G). In Pv interneurons, the peak amplitude, frequency, or decay of sEPSCs did not significantly differ between Pv-cKO and control mice (Figure 5H,I). In separate sets of experiments where both sEPSCs and sIPSCs were measured from Pv interneurons, at P42–56 or P152–192 when SWD frequencies were high, neither changes in sEPSC nor sIPSC properties were evident (Supplemental figure 6A–F). We also investigated whether loss of Cask resulted in changes of Pv interneuron intrinsic excitability and found that none of the resting membrane potential, rheobase current, action potential threshold, membrane resistance, or membrane capacitance was affected (Figure 5J,K).

Given the changes in sIPSCs but not uIPSCs onto pyramidal neurons in Pv-cKO mice, we next quantified the density of EYFP+ Pv interneurons in the S1 barrel cortex to determine if a reduction of cell number might contribute to this difference. Neither the overall density nor density in each cortical layer of EYFP+ neurons was reduced in Pv-cKO mice (Figure 5L,M).

Overall, *Cask* deficiency in Pv interneurons results in reduced sIPSC amplitude and frequency onto pyramidal neurons without altering the connectivity, action potential-evoked uIPSCs, postsynaptic properties, intrinsic excitability, or survival. Since majority of the sIPSCs recorded from pyramidal neuron somata originate from the perisomatic inhibitory synapses formed by Pv interneurons (Soltesz et al., 1995; Tremblay et al., 2016; Feldmeyer et al., 2018), a reduction of sIPSCs in Pv-cKO mice most likely reflects a reduction of spontaneous GABA release from Pv interneurons. Thus, these findings are consistent with the previous result that spontaneous, but not evoked, GABA release is reduced in cultured *Cask* KO neurons (Atasoy et al., 2007)).

### *Cask* deletion in Sst interneurons reduces evoked GABAergic transmission onto pyramidal neurons

We used a similar experimental design as described above to identify changes in Sst-cKO mice at P42–56 (Figure 6A,B), as no GTCSs have been observed yet at this age. While connectivity rate between layer 2/3 Sst interneurons and pyramidal neurons in the S1 did not differ between control (*Sst^Cre/+^;Rosa26^Ai3/+^*) and Sst-cKO (*Sst^Cre/+^;Cask^tm1c/y^;Rosa26^Ai3/+^;*) mice (Figure 6C), the first uIPSC amplitude was reduced by 59% in Sst-cKO mice (Figure 6D). Correspondingly, the short-term synaptic depression was less in Sst-cKO mice (Figure 6E), indicating that loss of Cask decreases the release probability of Sst interneurons and impairs inhibition onto pyramidal neurons. We also observed a small (8.9%) reduction of the peak amplitude of sIPSCs in pyramidal neurons, but the sEPSCs in either pyramidal neurons or Sst interneurons were not altered (Figure 6F–I). Additional experiments did not find changes in sEPSCs or sIPSCs in Sst interneurons either (Supplemental figure 6G–I). Furthermore, we found that the resting membrane potentials of Sst interneurons were significantly depolarized in Sst-cKO mice, but other intrinsic properties remained unchanged (Figure 6J,K). Finally, the overall density or density in each cortical layer of EYFP+ Sst interneurons was similar between Sst-cKO and control mice (Figure 6L,M).

**Figure 6:**
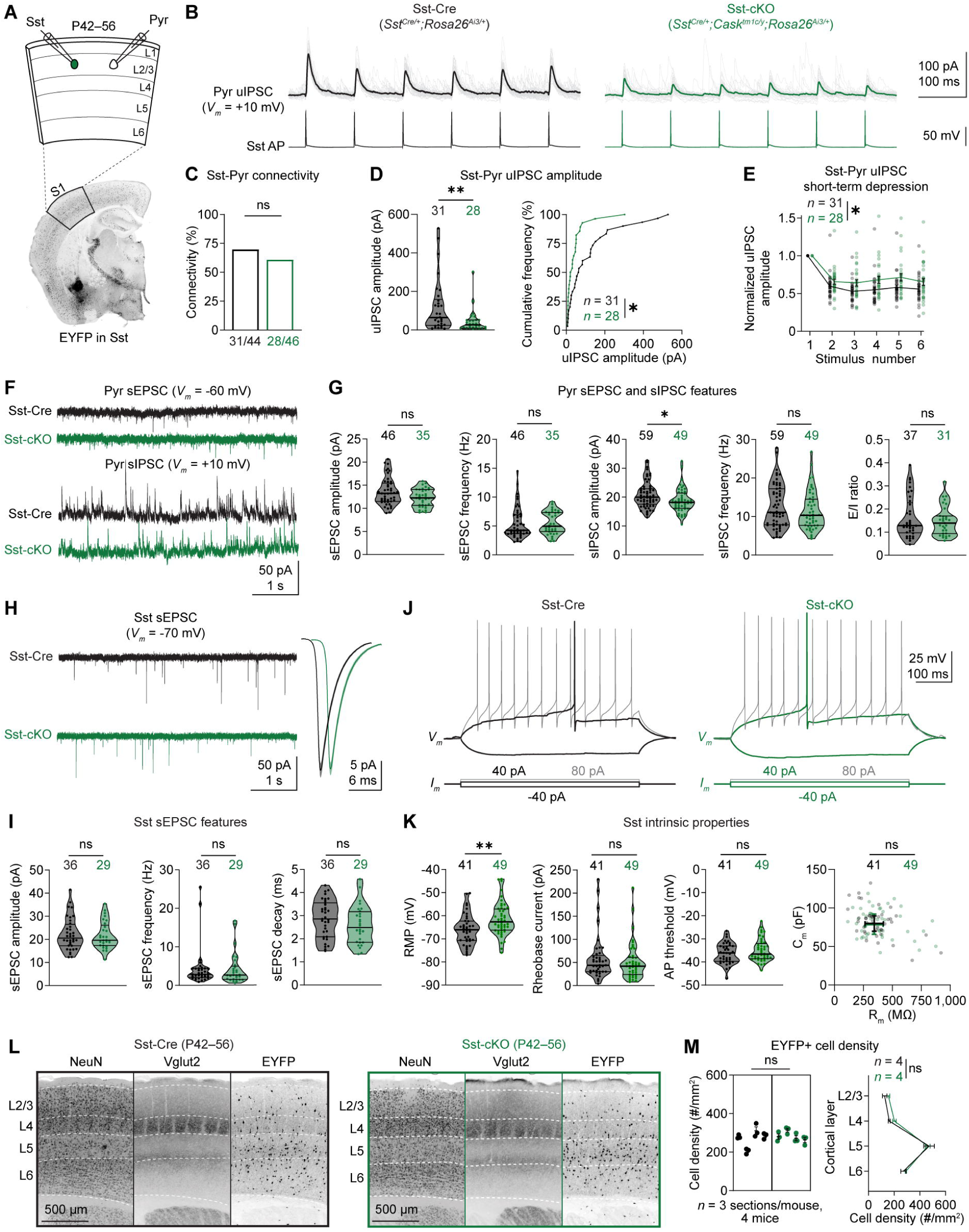
*Cask* deletion in Sst interneurons reduces unitary inhibition onto layer 2/3 pyramidal neurons. **(A)** Schematic of slice experiments (top), where a pair of adjacent (< 50 μm) Sst interneuron and pyramidal neuron were recorded in the layer 2/3 of S1 from 11 control Sst-Cre and 12 Sst-cKO male mice at P42–56. Sst interneurons were identified by the Cre-dependent expression of EYFP (bottom). **(B)** A train of six action potentials at 10 Hz in an Sst interneuron (lower panels) evoked uIPSCs in a pyramidal neuron held at +10 mV (upper panels). 30–50 individual traces (gray) and the average trace (color) are superimposed. **(C)** Unitary connection rates, Fisher’s exact test. **(D)** L Left, summary data of uIPSC amplitudes evoked by the first action potentials in the train. Lines indicate median (Sst-Cre, 64.27 pA; Sst-cKO, 26.39 pA) ± I.Q.R., Mann-Whitney rank test. Right, cumulative frequencies of uIPSC amplitude, Kolmogorov-Smirnov test. uIPSC amplitude was reduced in Sst-cKO mice. **(E)** uIPSC amplitudes during the train of action potentials were normalized by the amplitudes of the first uIPSC. Sst-cKO mice showed less synaptic depression. Data are mean ± S.E.M., two-way ANOVA. **(F)** Representative traces of sEPSCs (top) and sIPSCs (bottom) from pyramidal neurons. **(G)** Summary data of the sEPSC amplitude, frequency, sIPSC amplitude, frequency, and E/I ratio of pyramidal neurons. Lines indicate median ± I.Q.R., Mann-Whitney rank test. **(H)** Left, representative traces of sEPSCs from Sst interneurons. Right, the average sEPSC waveforms across all neurons from Sst-Cre (black) or Sst-cKO (purple) mice. **(I)** Summary data of sEPSC amplitude, frequency, and single-exponential decay time constant of Sst interneurons. Lines indicate median ± I.Q.R., Mann-Whitney rank test. **(J)** Membrane potentials (upper panels) in response to a hyperpolarizing (color), the rheobase (color), and a supra-rheobase (gray) current injections in Sst interneurons. **(K)** Summary data of resting membrane potential (RMP), rheobase current, action potential threshold, membrane capacitance (C_m_), and membrane resistance (R_m_) of Sst interneurons. Lines indicate median ± I.Q.R., Mann-Whitney rank test. **(L)** Representative fluorescence images of brain sections stained for NeuN, Vglut2, and EYFP from Sst-Cre or Sst-cKO mice. **(M)** Left, EYFP+ cell density of a 1000-μm-wide cortical column in S1 barrel cortex. Each symbol represents a section, and each column represents a mouse. Lines indicate mean (of sections) ± S.E.M., nested t-test. Right, EYFP+ cell density in each cortical layer, calculated as an average of the 3 sections. Data are mean (of mice) ± S.E.M., two-way ANOVA. Each symbol represents a synaptic connection or neuron, unless otherwise specified. The number (*n*) are indicated in the panels. ns, *P* > 0.05; *, *P* < 0.05; **, *P* < 0.01.

Altogether, loss of Cask in Sst interneurons results in a severe impairment of action potential-evoked uIPSCs and minor deficits in sIPSCs and resting membrane potential without altering their connectivity to pyramidal neurons, postsynaptic properties, or survival. The reduced evoked inhibition onto layer 2/3 pyramidal neurons is at least in part due to a decrease in release probability but could also results from a reduction of quantal amplitude. To test this possibility, we next isolated quantal release specifically from Sst interneurons in Sst-cKO mice.

### *Cask* deletion in Sst interneurons results in compensatory increases of quantal amplitude

We utilized an optogenetic protocol previously developed in our lab (Chen et al., 2020) to isolate quantal IPSCs (qIPSCs) mediated by the GABA release from specific neurons. By injecting a Cre-dependent adeno-associated virus (AAV) vector AAV9-EF1α-DIO-hChR2(H134R)-EYFP into the S1 region of Sst-Cre and Sst-cKO mice, we selectively expressed blue light-gated channelrhodopsin-2 (ChR2) in Sst interneurons. Between P42–56, we recorded miniature IPSCs (mIPSCs) from layer 2/3 pyramidal neurons of the S1 cortex in a modified ACSF containing tetrodotoxin (TTX), a voltage-gated sodium channel blocker, and Sr^2+^ instead of Ca^2+^. This modified ACSF enabled ramp-down blue light to stimulate ChR2-induced asynchronous release (Figure 7A,B). We then mathematically subtracted the mIPSCs recorded during the baseline period before blue light stimulation from those recorded during blue light stimulation to obtain the average amplitude, charge, and decay time constant of Sst interneurons-mediated qIPSCs. Loss of Cask in Sst interneurons caused a 23.3% and 21.8% increase in the quantal amplitude and charge, respectively, but did not affect the decay time constant (Figure 7C). This increase of qIPSC amplitude is probably a compensatory change in pyramidal neurons in response to the reduced inhibition. As uIPSC amplitude is determined by qIPSC amplitude, presynaptic release probability, and synaptic connection number, the increase of qIPSC amplitude together with the stark reduction of uIPSC amplitude suggest a strong reduction of presynaptic release probability since the connectivity from Sst interneurons to pyramidal neurons was unaltered (Figure 6C).

**Figure 7:**
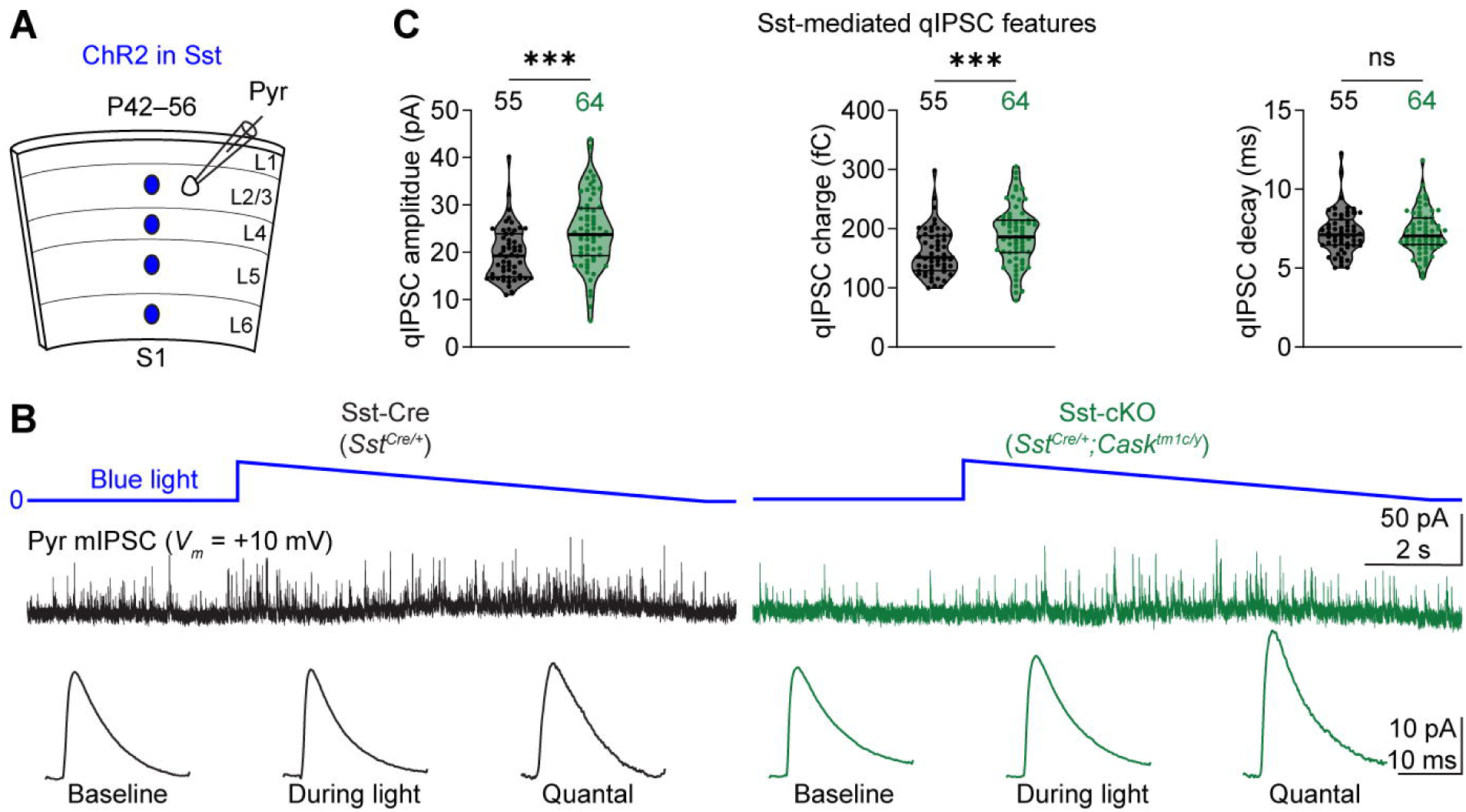
*Cask* deletion in Sst interneurons increases quantal inhibitory synaptic transmission. **(A)** Schematic of slice experiments, where ChR2 was expressed in Sst interneurons, and pyramidal neurons were recorded in the layer 2/3 of S1 from 5 control Sst-Cre and 5 Sst cKO male mice at P42–56. (**B**) Miniature IPSC (mIPSC) traces from pyramidal neurons held at +10 mV (middle panels). Blue light stimulation (top panels) activated ChR2 and increased mIPSC frequency. The quantal IPSC (qIPSC) trace was computed based on the average mIPSC trace during the baseline period and light stimulation period (bottom row). **(C)** Summary data of the average amplitude, charge, and single-exponential decay time constant of Sst interneuron-mediated qIPSCs in Sst-Cre and Sst-cKO mice. Lines indicate median ± I.Q.R., Mann-Whitney rank test. Sst-cKO mice show larger qIPSC average amplitudes and charge. Each symbol represents a neuron. The numbers (*n*) are indicated in the panels. ns, *P* > 0.05; ***, *P* < 0.001.

## DISCUSSION

Epilepsy is a cardinal feature of *CASK*-related disorder, but how *CASK* dysfunction disrupts neuronal circuits to produce distinct seizure phenotypes remains unclear. In this study, we established sex-specific mouse models of *CASK*-related disorder as bona fide models of epilepsy and used cell type-specific *Cask* deletion to identify inhibitory neurons as a critical cellular locus of epileptogenesis. Selective deletion of *Cask* in Pv or Sst interneurons revealed a striking divergence in seizure phenotypes that were characterized predominantly by spike-wave discharges (SWDs) or generalized tonic-clonic seizures (GTCSs), respectively. Electrophysiological analyses further uncovered distinct alterations in inhibitory synaptic transmission, including reduced spontaneous release associated with *Cask* deficiency in Pv interneurons and impaired action potential-evoked release associated with *Cask* deficiency in Sst interneurons. Together, these findings link the cellular functions of Cask in different cell types to distinct epileptic phenotypes and suggest that distinct modes of presynaptic dysfunction can engage different epileptic network dynamics.

Our study of both female heterozygous KO (Sox2-Het) and male hemizygous hypomorphic (Cask-hypo) mouse models identifies epilepsy as a key phenotype of *Cask* dysfunction *in vivo* and extends prior studies that primarily focused on microcephaly, cerebellar hypoplasia, and motor dysfunction (Najm et al., 2008; Srivastava et al., 2016; Patel et al., 2022; Guo et al., 2023; Mori et al., 2025). There has been a single report of a spontaneous hippocampal seizure in one female heterozygous KO mouse (Patel et al., 2020), but no studies have been conducted in the male model, even though males encompass nearly a third of humans with *CASK*-related disorder (Mori et al., 2023; Jiang et al., 2026a). Despite the underlying differences in residual Cask expression and cellular distribution between Sox2-Het and Cask-hypo mice (Figure 1N), the emergence of a shared epileptic phenotype of SWDs suggests a conserved mechanism of circuit dysfunction.

We addressed the cellular mechanisms of epilepsy with conditional deletion of *Cask* in different neuronal types by capitalizing on our novel *Cask* flox allele (*tm1c*), which does not affect Cask expression or exhibit any epileptic phenotype (Figure 1C–H, Figure 3D, Supplemental figure 3A). We pinpointed the loss of *Cask* in forebrain inhibitory neurons, but not cortical and hippocampal excitatory neurons, as critical drivers of epilepsy in male hemizygous mice. The phenotypic severity of Dlx5/6-cKO mice suggests that Cask function is particularly important for inhibitory neurons and network stability. Nevertheless, the absence of seizures in Vglut1-cKO and Emx1-cKO mice does not exclude contributions from excitatory neurons to disease pathophysiology, as deletion of *Cask* in all excitatory neurons (i.e., Vglut2-cKO) causes early lethality. One point of interest is the contrast between Cask-hypo mice and inhibitory neuron-specific male cKO mice. The former model only has SWDs, while several of the latter models have SWDs, GTCSs, other epileptic abnormalities, or combinations of these phenotypes. These differences suggest that the severity and nature of epilepsy depend on the degree of Cask dysfunction and vulnerability of certain cell types to Cask dysfunction. For instance, the approximately 70% Cask reduction in Cask-hypo mice may impact Pv interneurons more than Sst interneurons, thereby primarily causing SWDs, as observed in Pv-cKO mice. Future experiments testing if selectively restoring Cask levels in Pv interneurons can reduce SWDs in Cask-hypo mice will help evaluate this hypothesis.

A particular important finding of our study is the divergence of epileptic phenotypes between Pv-cKO and Sst-cKO mice, as it may be relevant to the diverse types of seizures observed in *CASK*-related disorder (Takanashi et al., 2012; Moog et al., 2015; Giacomini et al., 2021). Pv interneurons provide powerful and temporally precise inhibition of neuronal activity, including feedforward inhibition, whereas Sst interneurons regulate dendritic integration and contribute to feedback inhibition (Kepecs and Fishell, 2014). Reduction of feedforward inhibition due to dysfunction of Pv interneurons in cortical or thalamic circuits has been implicated in the SWDs of several rodent models (Paz et al., 2011; Panthi and Leitch, 2019; Abdelaal et al., 2022; Meyer and Maheshwari, 2024). This is likely the case in the Pv-cKO mice, though the progressive increase of SWDs in Pv-cKO mice may be contributed by the delayed loss of Cask because cortical Pv interneurons start expressing parvalbumin, and thus Cre, around P10–14 (del Rio et al., 1994; Taniguchi et al., 2011), and Cask has an estimated half-life of two weeks (Fornasiero et al., 2018). Dysfunction of Sst interneurons and feedback inhibition deficits have also been linked to generalized tonic-clonic seizures (Cossart et al., 2001; Ito-Ishida et al., 2015; Pothmann et al., 2019; Wengert et al., 2021; Drexel et al., 2022). These observations provide a plausible framework that different inhibitory circuits differentially contribute to the generation of epileptic activity, and *Cask* deletion in Pv or Sst interneurons disrupts different circuit operations, resulting in distinct seizure phenotypes. Loss of *Cask* in both cell types simultaneously, and thus dysfunction in multiple modes of inhibition, may explain why epilepsy in Dlx5/6-cKO mice is much more severe and has an earlier onset than Pv-cKO and Sst-cKO mice. Determining how different mechanisms of inhibition interact is an avenue of future investigation that may provide insights into how a shared genetic deficit can produce differences in seizure type, onset, and severity.

Our electrophysiological analyses provide further insight into the cellular consequences of *Cask* deletion and identify distinct presynaptic dysfunctions in Pv and Sst interneurons. Even though Cask has been implicated in regulating postsynaptic function and dendritic arborization in interneurons (Gao et al., 2018, 2019), we found no changes in the frequency or amplitude of synaptic inputs onto *Cask* deficient Pv or Sst interneurons, indicating that the postsynaptic function is largely preserved in these neurons. In Pv-cKO mice, the absence of detectable changes in the unitary connectivity from Pv interneurons to pyramidal neurons, unitary responses, intrinsic excitability, or Pv interneuron density argues again an impairment of evoked synaptic transmission or a gross loss of Pv interneurons. Instead, the reductions in sIPSC amplitude and frequency onto pyramidal neurons indicate a decrease in spontaneous GABA release from Pv interneurons, as among multiple classes of inhibitory neurons, Pv interneurons are the main source of sIPSCs in pyramidal neurons (Soltesz et al., 1995; Tremblay et al., 2016; Feldmeyer et al., 2018). This interpretation is consistent with previous results that spontaneous, but not evoked, release is reduced in *Cask* KO cultured cortical neurons (Atasoy et al., 2007). Thus, Cask primarily maintains spontaneous release at Pv interneuron synapses, and the effect of *Cask* deletion on evoked release may be more limited or were not detectable under our experimental conditions.

In contrast, Sst-cKO mice exhibit a significant decrease in the evoked uIPSC amplitude from Sst interneurons to pyramidal neurons, accompanied by less short-term synaptic depression and an increase of optogenetically-isolated qIPSC amplitude. These findings indicate that loss of *Cask* in Sst interneurons substantially reduces presynaptic release probability and uncover a previously unrecognized function of Cask in regulating evoked release. While the reduced uIPSC amplitude and increased qIPSC amplitude could also suggest fewer synaptic contacts from Sst interneuron onto pyramidal neurons, the preserved synaptic connectivity makes this possibility less likely. However, a change in the number or organization of synapses within individual connected pairs cannot be excluded and requires further studies.

The distinct presynaptic changes between Pv and Sst interneurons following *Cask* deletion suggest that Cask dysfunction have different functional consequences at these two types of inhibitory synapses and could be due to differences in their molecular composition and physiology. Although Cask is a shared presynaptic scaffolding protein, its interacting proteins and functional contributions may differ between these two types of synapses, such that its loss disrupts different components of the neurotransmitter release machinery. Furthermore, spontaneous and evoked release have shared but also distinct molecular processes (Guzikowski et al., 2026), and Cask may contribute differently to these processes at Pv and Sst interneuron synapses. These notions have been illustrated by the distinct effects of Synapsin-1 deletion on Pv and Sst interneuron synapses (Forte et al., 2020) ). Another example is that deletion of all three Neurexins, a key presynaptic binding partner of Cask, in cortical Sst, but not Pv, interneurons decreases evoked release via a reduction of action potential-evoked Ca^2+^ influx and presynaptic release probability (Jiang et al., 2026b).

Several inherent limitations of our study warrant considerations. First, while our electrophysiological analyses centered on inhibitory neurons in the somatosensory cortex, the Cre lines used in the study are not restricted to it. In particular, Pv- and Sst-expressing inhibitory neurons of the thalamic reticular nucleus may contribute to epilepsy (Paz et al., 2011; Hou et al., 2016; Clemente-Perez et al., 2017; Panthi and Leitch, 2019; Martinez-Garcia et al., 2020; Abdelaal et al., 2022; Meyer and Maheshwari, 2024). Second, the synaptic defects identified in Pv-cKO and Sst-cKO mice do not establish a direct causal relationship between a synaptic alteration and a specific seizure phenotype. Measurements of synaptic transmission and circuit activity *in vivo*, together with targeted rescue experiments, will be important for determining how these cellular alterations influence epileptic network dynamics. Third, we focused on male hemizygous cKO mice since *Cask* deletion was uniform in the neuronal type of interest. Whether similar inhibitory dysfunction mechanisms operate in female heterozygous mice, where there are mosaic populations, and how Cask dysfunction in non-neuronal populations contributes to epilepsy are important questions for future investigations. Nevertheless, our study reveals that epilepsy phenotypes and inhibitory synaptic impairments are not a uniform consequence of *Cask* loss, and Cask performs distinct functions in different types of neurons and synapses, highlighting the importance of linking genetic disruption to cell type-specific synaptic physiology and circuit function to understand how disruptions of a single neurodevelopmental disease gene can produce diverse neurological manifestations.

## METHODS

### Mice

A new *Cask* flox mouse line was generated from a *Cask* knock-out first allele (*tm1a*) that contains a trapping cassette flanked by two *FRT* sites and the exon 6 flanked by two *loxP* sites. The trapping cassette was removed by crossing to *Rosa26-Flpo* mice (Jackson Laboratory, JAX # 012930)(Raymond and Soriano, 2007), resulting in the *Cask* flox allele (*tm1c*). *Cask* flox mice were genotyped by PCR using the forward primer 5′-GTATATCTGCACTTGTAGAATACC-3′ and a pair of reverse primers 5′-TTGGAAATTGGTTTGGCAAGATGG-3′ for wildtype (WT; 461 bp) and 5’-TGAACTGATGGCGAGCTCAGACC-3’ for *tm1c* (616 bp) alleles. *Cask* flox mice were maintained on the C57BL/6J background by crossing to WT C57BL/6J mice (JAX # 000664).

Female heterozygous *Cask* flox mice (*Cask^tm1c/+^*) were crossed to male hemizygous C57BL/6J-congenic *Sox2-Cre* (JAX # 008454) (Hayashi et al., 2002) mice to generate female offspring: WT, *Cask^tm1c/+^, Sox2-Cre^Tg^*^/+^, and *Sox2-Cre^Tg^*^/+^;*Cask^tm1c/+^*and male offspring: WT, *Cask^tm1c/y^, Sox2-Cre^Tg^*^/+^, and *Sox2-Cre^Tg^*^/+^;*Cask^tm1c/y^*. Female heterozygous *Cask^tm1sud/+^* mice (JAX # 006382)(Atasoy et al., 2007) were crossed with male WT C57BL/6J to generate male offspring: WT and *Cask^tm1Sud/y^*.

Male heterozygous C57BL/6J-congenic *Viaat-ires-Cre* mice (JAX # 028862) (Vong et al., 2011) were crossed with female *Cask^tm1c/+^*mice to generate female and male offspring: WT, *Cask^tm1c/+^* or *Cask^tm1c/y^*, *Viaat^Cre/+^*, and *Viaat^Cre/+^*;*Cask^tm1c/+^*or *Viaat^Cre/+^*;*Cask^tm1c/y^* mice. Male heterozygous C57BL/6J-congenic *Vglut2-ires-Cre* mice (JAX # 028863)(Vong et al., 2011) or hemizygous *Dlx5/6-Cre* mice (JAX # 028863)(Monory et al., 2006) on a C57BL/6J background were crossed with female *Cask^tm1c/+^*mice similarly. A subset of *Dlx5/6-Cre^Tg/+^*;*Cask^tm1c/^y* mice were generated by crossing male *Dlx5/6-Cre^Tg/+^* mice with female homozygous *Cask^tm1c/tm1c^* mice. Male homozygous C57BL/6J-congenic *Emx1-ires-Cre* (JAX # 005628) (Gorski et al., 2002) were crossed to female *Cask^tm1c/+^* mice to yield *Emx1^Cre/+^* and *Emx1^Cre/+^;Cask^tm1c/y^*. Male homozygous C57BL/6J-congenic *Vglut1-ires-Cre-D* mice (JAX # 037512)(Harris et al., 2014), C57BL/6J-congenic *Pv-ires-Cre* mice (JAX # 017320) (Hippenmeyer et al., 2005), and *Sst-ires-Cre* mice (JAX # 013044)(Taniguchi et al., 2011) on a C57BL/6;129S4 background were crossed with female *Cask^tm1c/+^* mice similarly. Male hemizygous *Htr3a-Cre mice* (NO152 line, MMRRC # 036680-UCD)(Gerfen et al., 2013; Miyoshi et al., 2015) on a FVB;C57BL6;129;Swiss;CD1 background were crossed to female *Cask^tm1c/+^* mice to yield male WT, *Cask^tm1c/y^*, *Htr3a-Cre^Tg/+^*, and *Htr3a-_CreTg/+;Casktm1c/y_*.

Male homozygous C57BL/6J-congenic *Rosa26-CAG-LSL-EYFP* mice (Ai3, JAX # 007903) were crossed to female *Cask^tm1c/+^* mice, and the male offspring *Cask^tm1c/y^;Rosa26^Ai3/+^*mice were crossed to female *Rosa26^Ai3/Ai3^* to yield *Cask^tm1c/+^;Rosa26^Ai3/Ai3^*female mice. Male homozygous *Pv-ires-Cre* mice were crossed with females *Cask^tm1c/+^;Rosa26^Ai3/Ai3^* to generate male *Pv^Cre/+^;Rosa26^Ai3/+^*and *Pv^Cre/+^;Cask^tm1c/y^*;*Rosa26^Ai3/+^*mice for slice electrophysiology experiments. Male homozygous *Sst-ires-Cre* mice were crossed with female *Cask^tm1c/+^;Rosa26^Ai3/Ai3^*mice similarly.

Mice were housed in an Association for Assessment and Accreditation of Laboratory Animal Care International-certified animal facility on a 14 hr light/10 hr dark cycle. All procedures to maintain and use mice were performed in strict accordance with the recommendations in the Guide for the Care and Use of Laboratory Animals of the National Institutes of Health and were approved by the Institutional Animal Care and Use Committee at Baylor College of Medicine (protocol AN-6544).

### Western blot and RT-qPCR

Male mice at postnatal day 0 (P0) were anesthetized and decapitated. Brains were hemisected without further dissection. Juvenile female mice at 1 month of age and adult male mice at 2 months of age were anesthetized and decapitated. After harvesting the brains, different brain regions were dissected from either the right or left hemisphere. All tissues were frozen at −80°C until analysis.

Western blots were performed as described previously with modifications (Chen et al., 2020; Lee et al., 2023). Briefly, protein was extracted in a lysis buffer containing 50 mM Tris-HCl (pH 7.6), 150 mM NaCl, 1 mM EDTA, 1% Triton X-100, 0.5% Na-deoxycholate, 0.1% SDS, and Xpert Proteinase Inhibitor Cocktail Solution (GenDepot, catalog # P3100). 30 µg of protein from the P0 male mice and juvenile female mice or 15 µg of protein from the adult male mice were loaded. Cask and the control protein β3-Tubulin were detected by a mouse anti-Cask (epitope amino acids 318–415, BioLegend, catalog # 830601, lot # B420734, 1:1000) or rabbit anti-Cask (epitope N-terminus within the first 143 amino acids, Cell Signaling, catalog # 2878S, 1:1000 dilution) and a mouse anti-β3-Tubulin (Proteintech, catalog # 66240-1-Ig, lot # 10025689, 1:100,000 dilution). Primary antibodies were identified by a goat anti-mouse IRDye 800LT antibody (LI-COR Biosciences, catalog # 925–33120, lot # D60303-02, 1:10,000) or a goat anti-rabbit IRDye (LI-COR Biosciences, catalog # 925-68071, lot # D41015-06, 1:10,000). Proteins were visualized and quantified using an Odyssey CLx Imager and Image Studio Lite 6.1 (LI-COR Biosciences). The expression levels of Cask were normalized to β3-Tubulin for individual samples, then normalized to the average normalized Cask levels of the WT mice from the sample blots.

For RT-qPCR, RNA was extracted using Invitrogen PureLink^TM^ RNA Mini Kit (ThermoFisher, catalog # 12183025). 300 μl of lysis buffer and 0.1% (vol/vol) of β-mercaptoethanol were added to 5–10 mg of the brain tissues. Homogenization was performed for 1 min at 4 m/s using a bead mill homogenizer (Bead Raptor Elite, Omni International). Samples were treated with DNase I (Sigma-Aldrich, Catalog # AMPD1), and RNA concentrations were measured. Reverse transcription was done with High-Capacity cDNA Reverse Transcription Kit (ThermoFisher, catalog # 4368814). qPCR reaction was performed in triplicate for each sample using amfiSure qGreen Q-PCR Master Mix (GenDepot, catalog # Q5600). *Cask* mRNA was detected via the forward primer: 5’-ACCCATGGCTTAAGGAGCG-3’ (spanning exon 8–9) and reverse primer: 5’-CCTGAGGAGGTAGGGTCTTC-3’ (spanning exon 10–11). *Gapdh* mRNA was detected via the forward primer: 5’-GGAGAGTGTTTCCTCGTCCC-3’ and reverse primer: 5’-ATGAAGGGGTCGTTGATGGC-3’. The C_q_ values of *Cask* and *Gapdh* for each sample was determined by averaging across the technical triplicates. The ΔC_q_ was calculated by subtracting the *Gapdh* C_q_ from the *Cask* C_q_. The ΔΔC_q_ for each sample was then obtained by subtracting the average ΔC_q_ of the WT group. The relative *Cask* expression for each sample was calculated by raising 2 to the power of the negative ΔΔC_q_ (2^-ΔΔCq^) and normalized to the average of WT group.

### Video-EEG/EMG recording and seizure detection

Video-electroencephalography and electromyography (EEG/EMG) recordings were performed as previously described with minor modifications (Chen et al., 2020). Briefly, mice between 3–6 weeks of age were anesthetized with 1.25–2.5% isoflurane in oxygen. Craniotomies (approximate diameter of 0.25 mm) were performed with a round bur (0.25 mm diameter) at the coordinates below with distances relative to the bregma and normalized by the distance between bregma and lambda (BL). Perfluoroalkoxy polymer (PFA)-coated silver wire electrodes (A-M Systems, catalog # 786000, 127 μm bare diameter, 177.8 μm coated diameter) were used for grounding at the right frontal cortex and recording at the left frontal cortex (anterior posterior (AP), 0.42 of BL; medial lateral (ML), 0.356 of BL; dorsal ventral (DV), −1.5 mm); left and right somatosensory cortices (AP, −0.34 of BL; ML, ±0.653 of BL; DV, −1.5 mm); and referencing at the cerebellum. An EMG recording and an EMG reference electrode were inserted into the neck muscles. Mice were given 3 days to 1 week to recover from the surgeries. Before recording, mice were individually habituated in the recording chambers overnight. EEG/EMG signals (5,000 Hz sampling rate with a 0.5 Hz high-pass filter) and videos (30 frames per second) were recorded synchronously for continuous 48–72 hours using a four-channel EEG/EMG tethered system (Pinnacle Technology).

Spike-wave discharges (SWDs) were identified by generating putative candidates with custom-written code in MATLAB (MathWorks) followed by the classification of candidates with a convolutional neural network in Python that has been trained with manually labeled EEG segments (Chen et al., 2020). Identified SWDs were confirmed by visual inspection. All video-EEG/EMG recordings were manually reviewed offline by trained individuals to identify generalized tonic-clonic seizures and other epileptic events. Epileptic events lasting more than 5 seconds, with or without clinical convulsions, were manually annotated for the start and end and then graded by the highest level reached on the modified Racine scale (Van Erum et al., 2019). Handling-induced seizures were qualitatively recorded by animal husbandry or individual researchers if an animal was noted to exhibit a generalized convulsive seizure during routine animal handling or cage-changes.

### AAV injection

Recombinant AAV vector AAV9-EF1α-DIO-hChR2(H134R)-EYFP (Addgene # 20298) was obtained from the Penn Vector Core (catalog # AV-9-20298P). To express ChR2 in Pv or Sst interneurons, 200 nL of AAV9-EF1α-DIO-hChR2(H134R)-EYFP (1.46×10^12^ genome copies/mL) was injected into the somatosensory cortex of *Pv^Cre/+^* and *Pv^Cre/+^;Cask^tm1c/y^*, or *Sst^Cre/+^ and Sst^Cre/+^;Cask^tm1c/y^* mice, respectively, at P0–1 as previously described (Xue et al., 2014; Messier et al., 2018) with an UltraMicroPump III and a Micro4 controller (World Precision Instruments).

### Brain slice electrophysiology

Mice were anesthetized by an intraperitoneal injection of a ketamine and xylazine mix (240 mg/kg and 48 mg/kg, respectively) and transcardially perfused with cold (0–4 C) slice cutting solution containing 80 mM NaCl, 2.5 mM KCl, 1.3 mM NaH_2_PO_4_, 26 mM NaHCO_3_, 4 mM MgCl_2_, 0.5 mM CaCl_2_, 20 mM D-glucose, 75 mM sucrose, and 0.5 mM sodium ascorbate (315 mosmol, pH 7.4, saturated with 95% O_2_/5% CO_2_). Brains were removed, hemi-sected, and then sectioned in the cutting solution with a VT1200S vibratome (Leica) to obtain 300 μm coronal slices. Slices containing the primary somatosensory cortex were collected and incubated in a custom-made interface holding chamber in which a cellulose nitrate filter (Sartorius, catalog # 13006--50----ACR) was placed on the top of a 50 × 35 mm round crystallizing dish (Chemglass Life Sciences) filled entirely with slice cutting solution that was saturated with 95% O_2_/5% CO_2_. The dish was placed in a larger 70 × 50 mm round crystallizing dish (Pyrex) filled with approximately 1 cm of slice cutting solution that was saturated with 95% O_2_/5% CO_2_. The holding chamber was covered by a cap lined with filter paper to ensure humidity. Slices were incubated on the cellulose nitrate filter at 34 C for 30 min and then at room temperature for 20 minutes to 7 hours until they were transferred to the recording chamber.

Recordings were performed on slices submerged in artificial cerebrospinal fluid (ACSF) containing 119 mM NaCl, 2.5 mM KCl, 1.3 mM NaH_2_PO_4_, 26 mM NaHCO_3_, 1.3 mM MgCl_2_, 2.5 mM CaCl_2_, 20 mM D-glucose, and 0.5 mM sodium ascorbate (305 mosmol, pH 7.4, saturated with 95% O_2_/5% CO_2_, perfused at 3 mL/min) at 30–32 C. Whole-cell patch clamp recordings were performed using a K^+^-based pipette solution containing 142 mM K^+^-gluconate, 10 mM HEPES, 1 mM EGTA, 2.5 mM MgCl_2_, 4 mM ATP-Mg, 0.3 mM GTP-Na, and 10 mM Na_2_-phosphocreatine (295 mosmol, pH 7.35) or a Cs^+^-based pipette solution containing 121 mM Cs^+^-methanesulfonate, 10 mM HEPES, 10 mM EGTA, 1.5 mM MgCl_2_, 4 mM ATP-Mg, 0.3 mM GTP-Na, 10 mM Na_2_-phosphocreatine, and 2 mM QX314-Cl (295 mosmol, pH 7.35). Reported membrane potentials were not corrected for liquid junction potentials that were experimentally measured as 12.5 mV for the K^+^-based pipette solution and 9.5 mV for the Cs^+^-based pipette solution (Messier et al., 2018).

To identify the primary somatosensory cortex and barrel cortex, slices were visualized with video-assisted infrared differential interference contrast imaging (DIC) via an air objective (4× or 5×, 0.1 numerical aperture) on an upright SliceScope Pro 1000 microscope (Scientifica) and an infrared IR-1000 CCD camera (DAGE-MTI). Neurons were then visualized with DIC, and fluorescent neurons were identified by epifluorescence imaging via a water immersion objective (40×, 0.8 numerical aperture). Pyramidal neurons were identified based on their morphology, and Pv and Sst interneurons were identified by the Cre-dependent expression of EYFP. For photostimulation of ChR2-expressing neurons, blue light was emitted from a collimated light-emitting diode (LED) of 455 nm. The LED was driven by an LED driver (Mightex) under the control of an Axon Digidata 1550 Data Acquisition System and Clampex 10.7. Light was delivered through the reflected light fluorescence illuminator port and the 40× objective. Data were acquired at 10 kHz and low-pass filtered at 4 kHz with an Axon Multiclamp 700B amplifier and an Axon Digidata 1550 or 1440 Data Acquisition System under the control of Clampex 10.7 (Molecular Devices) and analyzed offline using AxoGraph 1.7.6 (AxoGraph Scientific) or custom-written code in MATLAB (MathWorks). For voltage clamp experiments, only recordings with series resistance below 20 MΩ were included in synaptic current measurements.

### Paired whole-cell recordings in pyramidal neurons and interneurons

Pairs of pyramidal neuron and interneuron in layer 2/3, approximately 200–300 μm from the pia, that were less than 50 μm apart by the Euclidean distance were identified and patched. A pyramidal neuron was patched first with the Cs^+^-based patch pipette solution, followed by a Pv or Sst interneuron with the K^+^-based pipette solution. Immediately after breaking into the interneuron, neuronal intrinsic excitability was examined. The resting membrane potential was recorded in the whole-cell current clamp mode within the first minute after break-in. After bridge balance, the membrane resistance and capacitance were measured by injecting a 500 ms hyperpolarizing current pulse of 60 or 40 pA (for Pv or Sst interneurons, respectively) to generate a small membrane potential hyperpolarization (5–10 mV) from the resting membrane potential. After initially finding an approximate rheobase current, depolarizing currents were increased in 5- or 2-pA steps (for Pv or Sst interneurons, respectively) to identify the rheobase currents. Following this, pyramidal neurons and interneurons were recorded in whole-cell voltage clamp mode at the reversal potential for inhibition (pyramidal neuron, -60 mV; Pv or Sst interneurons, -70 mV) to record spontaneous excitatory postsynaptic currents (sEPSCs) for 300–500 seconds. Subsequently, pyramidal neurons were voltage clamped at the reversal potential for excitation (+10 mV) and Pv or Sst interneurons were switched back to whole-cell current clamp mode. Action potentials were elicited in Pv or Sst interneurons by a train of 6 depolarizing current steps (2 msec, 3 nA) at 10 Hz with 15 s intervals between sweeps for 30–50 sweeps. To determine unitary inhibitory postsynaptic currents (uIPSCs), all sweeps were averaged, with some excluded if large disruptive spontaneous IPSCs (sIPSCs) were present, and uIPSC peaks were measured as the maximal currents in the 10 msec following the end of the stimulation, subtracting the average of the 10-msec baseline prior to stimulation. The uIPSC amplitude was defined as the peak of the first uIPSC. A Pv or Sst interneuron was considered to connect with a pyramidal neuron when the uIPSC amplitude was at least five times the baseline standard deviation. The following five uIPSC peaks were normalized to the first uIPSC peak to determine short-term synaptic depression. During each sweep, sIPSCs were also recorded in the pyramidal neurons for 12.5 seconds from 1.25 seconds after the 6th stimulus to the end of the sweep.

### Cell type-specific quantal IPSCs in pyramidal neurons

To isolate Pv or Sst interneurons-mediated quantal IPSCs, pyramidal neurons were recorded with the Cs^+^-based patch pipette solution in whole-cell voltage clamp mode at the reversal potential for excitation (+10 mV). A modified ACSF containing 4 mM MgCl_2_ and 0.5 mM SrCl_2_ without CaCl_2_, TTX (1 μM), NBQX (10 μM), and CPP (10 μM) were used to desynchronize synaptic vesicle fusion, block synaptic excitation, and reduce overall activity in the slices. Typically, 10–20 sweeps were recorded for each neuron with 40 s intervals between sweeps. During each sweep, miniature IPSCs (mIPSCs) were recorded during the 10-s baseline period and the 10-s blue light stimulation period. The light intensity (0.20–47.46 mW/mm^2^) was ramped down to reduce the tonic currents. The average amplitude, charge, and decay time constant of quantal IPSCs (qIPSCs) released from ChR2-expressing interneurons were computed as *Z_quantal_* = (*Z_light_* × *f_light_* - *Z_baseline_ × f_baseline_*)/ (*f_light_* - *f_baseline_*), where *Z* is the average amplitude, charge, or decay time constant of mIPSCs during the given period, and *f* is the frequency of mIPSCs during the given period. The average traces of qIPSCs were computed similarly using the average traces of mIPSCs from the light stimulation period and baseline period. Data were excluded if the number of qIPSCs was less than 150 or 10% of the mIPSCs during the baseline period. The criterion of at least 150 qIPSCs was chosen because the likelihood to estimate the parameters with less than 10% error from 150 events is higher than 95%.

### sEPSCs and sIPSCs in interneurons

Pv and Sst interneurons in layer 2/3, approximately 200–300 μm from the pia, were identified by the Cre-dependent expression of EYFP and recorded with the Cs^+^-based patch pipette solution in the whole-cell voltage clamp mode. sEPSCs were recorded at the reversal potential for inhibition (-60 mV) for 300–900 seconds and sIPSCs at the reversal potential for excitation (+10 mV) for 300–900 seconds.

### sEPSCs, sIPSCs, and mIPSCs detection

Data were digitally low-pass filtered at 2 kHz offline. sEPSCs, sIPSCs, and mIPSCs events were detected by a scaled-template algorithm (AxoGraph 1.7.6) using the following templates and parameters. Pyramidal neuron sEPSCs: template length, 10 ms; baseline, 3 ms; amplitude, –2 pA; rise time, 0.6 ms; decay time, 3 ms; and threshold, 3× standard deviation (SD) above baseline. Pyramidal neuron sIPSCs and mIPSCs: template length, 20 ms; baseline, 3 ms; amplitude, 2 pA; rise time, 0.6 ms; decay time, 10 ms; and threshold, 2.5× SD above baseline. Pv interneuron sEPSCs: template length, 5 ms; baseline, 1.5 ms; amplitude, –2 pA; rise time, 0.2 ms; decay time, 1 ms; and threshold, 3.25× SD above baseline. Pv interneuron sIPSCs: template length, 10 ms; baseline, 3 ms; amplitude, 2 pA; rise time, 0.27 ms; decay time, 3.7 ms; and threshold, 3 SD× above baseline. Sst interneuron sEPSCs: template length, 10 ms; baseline, 3 ms; amplitude, –2 pA; rise time, 0.4 ms; decay time, 1.9 ms; and threshold, 3.25 SD× above baseline. Sst interneuron sIPSCs: template length, 20 ms; baseline, 3 ms; amplitude, 2 pA; rise time, 0.6 ms; decay time, 13 ms; and threshold, 3.3× SD above baseline.

### Immunofluorescence staining and microscopy

To quantify the densities of Pv and Sst interneurons labeled by the Cre-dependent expression of EYFP, the brain hemispheres not used for slice electrophysiology were fixed overnight (10–12 hours) in 4% paraformaldehyde at 4 C, cryoprotected with 30% sucrose, and sectioned into 50 μm coronal slices using a Leica CM3050 S Cryostat. Sections containing the primary somatosensory cortex (S1) were incubated in blocking solution (0.2% Triton X-100 in PBS with 10% normal goat serum) for 1 hour and then with primary antibodies for 24 hours at 4 C. Primary antibodies were diluted in the blocking solution: guinea pig anti-NeuN (MilliporeSigma catalog # ABN90, lot # 4266748, 1:1000), chicken anti-GFP (Abcam, catalog # ab13970, lot # 1018753-19, 1:1000), and rabbit anti-Vglut2 (Synaptic Systems, catalog # 135 403, lot # 4-82, 1:1000). Sections were washed in 0.2% Triton X-100 in PBS and then incubated with the following secondary antibodies diluted 1:1000 in the blocking solution for 24 hours at 4 C: goat anti-guinea pig IgG (H+L) conjugated with DyLight 405 (Invitrogen, catalog # C840H41, lot # 168171, 1:200), goat anti-chicken IgG (H+L) conjugated with Alexa Fluor 488 (Invitrogen, catalog # A-11039, lot # 2180688, 1:1000), and goat anti-rabbit IgG (H +L) conjugated with Alexa Flour 555 (Invitrogen, catalog # A-21429, lot # 1622776, 1:1000). Sections were washed in 0.2% Triton X-100 in PBS and mounted in ProLong Diamond Antifade Mountant (Thermo Fisher Scientific, catalog # P36961, lot # 3225000). The images of whole coronal sections were acquired at a low magnification (20×), and the barrel cortices identified by the Vglut2 signal were acquired at a higher magnification (63×) on an Axio Zoom.V16 Fluorescence Stereo Zoom Microscope (Zeiss). Three brain sections were imaged and quantified per mouse. Images were visualized in ImageJ (National Institutes of Health), and a 1000-μm wide region of interest (ROI) was cropped across the entire cortical depth. Additional ROIs for each of cortical layers 2–6 was drawn based on the size and density of neurons labeled by the NeuN signal. Within each region, the number of EYFP+ cells were counted. Cell density was quantified by dividing the number of cells over the area of cortical layers 2–6 or each layer individually.

To visualize the overall brain morphology, mice were anesthetized by an intraperitoneal injection of a ketamine and xylazine mix (240 mg/kg and 48 mg/kg, respectively) and transcardially perfused with PBS, followed by 4% paraformaldehyde in PBS (pH 7.4). Brains were then post-fixed, cryoprotected, and sectioned into 50 μm sagittal slices, and stained with guinea pig anti-NeuN (MilliporeSigma catalog # ABN90, lot # 4266748, 1:1000) and goat anti-guinea pig IgG (H+L) conjugated with DyLight 405 (Invitrogen, catalog # C840H41, lot # 168171, 1:200) as described above. The images of whole sagittal sections were acquired with a MICA Microhub (Leica) at low magnification (10×) via tile scanning.

### Experimental design and statistical analysis

All efforts were made to use age-matched littermate controls for all experiments. Reported sample numbers (*n*) represent independent biological replicates, unless otherwise specified, that are the numbers of tested mice or recorded neurons. Statistical analyses were performed with Prism 11 (GraphPad Software). Binomial or Chi-square test was used for comparison of parts-of-a-whole data, Fisher’s exact test for contingency data, Log-rank (Mantel-Cox) test for survival data, and two-Way ANOVA or a mixed-effect model for comparisons across multiple timepoints. Otherwise, statistical tests that do not assume a Gaussian distribution and non-parametric tests were used. Statistical tests were two-tailed with an alpha of 0.05. The details of all statistical tests, numbers of replicates, and *P* values are reported in Supplementary file 1.

## ACKNOWLEDGEMENT

We thank Eli, Debra, and Steven Sukin for inspiring this work, Baylor College of Medicine Genetically Engineered Rodent Models Core led by Dr. Jason Heaney for the ES cell work and blastocyst injection to generate *Cask* knock-out first (*tm1a*) mice, Dr. Atul Maheshwari for discussion, and Dr. Wu Chen for technical support. We acknowledge the use of ChatGPT and Gemini to assist in editing the language of the manuscript. We reviewed and edited the content for accuracy and take full responsibility for the final text.

This work was supported in part by the Main Street America Fund and CASK Research Fund at Texas Children’s Hospital, the Eunice Kennedy Shriver National Institute of Child Health and Human Development (P50HD103555 to Baylor College of Medicine Intellectual and Developmental Disabilities Research Center, Neurovisualization Core; F30HD114410 to K.J.), the National Cancer Institute (P30CA125123 to Baylor College of Medicine Cancer Center). M.X. is a Caroline DeLuca Scholar.

## SUPPLEMENTAL FIGURE LEGENDS

**Supplemental figure 1: Reduced body weights and microcephaly in sex-specific mouse models of *CASK*-related disorder.**

**(A)** Pre-weaning body weights at P10 and P21 were reduced in Sox2-Het and Cask-hypo mice compared to controls. Data are mean ± S.E.M., two-way ANOVA. **(B)** Sox2-Het mice continued to show mild, but not significant body weight reductions as they aged while Cask-hypo mice continued to have significantly reduced body weights. Data are mean ± S.E.M., two-way ANOVA. **(C)** During surgeries for implanting EEG/EMG electrodes at P35–42 the measured Bregma-Lambda (BL) distance was significantly smaller in Sox2-Het and Cask-hypo mice compared to controls, indicating microcephaly at this age. Data are median ± I.Q.R., Mann-Whitney test. **(D)** Brain weight measured after the final video-EEG/EMG recordings was significantly reduced in Sox2-Het and Cask-hypo mice. Data are mean ± S.E.M., unpaired t-test. Filled (male) and open (female) symbols represent individual mice. The numbers (*n*) are indicated in the panels. ns, *P* > 0.05; *, *P* < 0.05; **, *P* < 0.01; ***, *P* < 0.001; ****, *P* < 0.0001. **(G,H)** Representative fluorescence images from mid-line sagittal brain sections stained with NeuN. Note the global microcephaly and the marked cerebellar hypoplasia in Sox2-Het (G) and Cask-hypo (H) mice.

**Supplemental figure 2: *Cask* deletion in broad inhibitory or excitatory neurons causes early lethality.**

**(A)** Offspring from *Viaat^Cre/^*^+^ (top) or *Vglut2^Cre/+^* (bottom) crossed to *Cask^tm1c/+^*. The color scheme is maintained across all panels for male and female mice separately. **(B)** At P10, the numbers of male offspring were less than the Mendelian expectation for *Viaat^Cre/^*^+^ (top) and *Vglut2^Cre/+^*(bottom) crosses, suggesting fewer males were born or survived to P10. Binomial test. **(C)** Pie charts representing the observed numbers of offspring with different genotypes at P10–14. Female offspring showed the expected Mendelian ratios, but male offspring Viaat-cKO and Vglut2-cKO mice were nearly absent. Chi-square test. **(D)** Viaat-cKO and Vglut2-cKO mice that were monitored from P0 showed significantly reduced survival compared to male control mice. WT, Flox, and Cre control mice were combined from both crosses. Log-rank (Mantel-Cox) test with Holm-Šídák’s multiple comparisons test. Most Viaat-cKO and Vglut2-cKO mice died at P0–1. Two Viaat-cKO mice that survived to P14 were observed to exhibit convulsive-like events. The numbers (*n*) are indicated in the panels. ns, *P* > 0.05; **, *P* < 0.01; ****, *P* < 0.0001.

**Supplemental figure 3: Spike-wave discharge frequency is not elevated in *Cask^tm1c/y^* (Flox), Vglut1-cKO, or Emx1-cKO mice.**

**(A,B)** Summary data of SWD frequency of *Cask^tm1c/y^*(Flox) and WT male mice (A) or Vglut1-cKO, Emx1-cKO, and their control mice (B). Data are mean ± S.E.M., two-way ANOVA. Symbols represent individual mice. The numbers (*n*) are indicated in the panels. ns, *P* > 0.05.

**Supplemental figure 4: Additional epileptic events and EEG abnormalities in Dlx5/6-cKO mice.**

**(A)** EEG/EMG traces of a Dlx5/6-Cre male mouse during NREM sleep. **(B)** EEG/EMG traces of a Dlx5/6-cKO mouse showing EEG abnormalities associated with NREM sleep including continuous slow spike-waves and poly-spikes/ inter-ictal spikes followed by EEG attenuation. **(C)** EEG/EMG traces of a Dlx5/6-Cre male mouse during active (left) and quiet (right) wake state. **(D)** EEG/EMG traces from a Dlx5/6-cKO mouse showing a cluster of myoclonic seizure. A high voltage EEG spike was associated with jerking of the neck or body. **(E,F)** EEG/EMG traces of two Dlx5/6-cKO mice showing clusters of spike-wave discharges (SWDs) at a typical 6–8 Hz frequency indicated by arrow heads as well as a period of slower frequency spike-waves (pink line).

**Supplemental figure 5: Additional EEG abnormalities in Pv-cKO mice.**

**(A)** EEG/EMG traces of a Pv-Cre male mouse during NREM sleep. Expanded sections highlight the slow waves. **(B)** EEG/EMG traces of a Pv-cKO mouse during NREM sleep. The slow oscillations were higher in amplitude, and there were occasional inter-ictal spikes.

**Supplemental figure 6: *Cask* deletion in Pv or Sst interneurons does not alter their synaptic inputs.**

**(A)** Schematic of slice experiments, where Pv interneurons were recorded in the layer 2/3 of the primary somatosensory cortex (S1) from 7 Pv-Cre and 6 Pv-cKO male mice at P42–P56. At this age Pv-cKO mice had low frequency of SWDs. **(B)** Representative traces of sEPSCs (left panels) and sIPSCs (right panels) from Pv interneurons. **(C)** Summary data of the sEPSC amplitude, frequency, sIPSC amplitude, frequency, and E/I ratio of Pv interneurons from Pv-Cre (black) and Pv-cKO (purple) mice. Lines indicate median ± I.Q.R., Mann-Whitney rank test. **(D–F)** Same as in (A-C), but for 6 Pv-Cre and 7 Pv-cKO mice at P152–192. At this age, Pv-cKO mice exhibited high frequency of SWDs. **(G–I)** Same as in (A-C), but for layer 2/3 Sst interneurons in 5 Sst-Cre (black) and 5 Sst-cKO (green) mice at P42–P56. At this age, Sst-cKO mice did not exhibit GTCSs. Each symbol represents a neuron. The numbers (*n*) are indicated in the panels. ns, *P* > 0.05.

