## Supplementary figures and images for "*Cask* dysfunction in inhibitory neurons reveals distinct synaptic and epileptic phenotypes in mouse models of *CASK*-related disorder"

### Supplemental figure 1

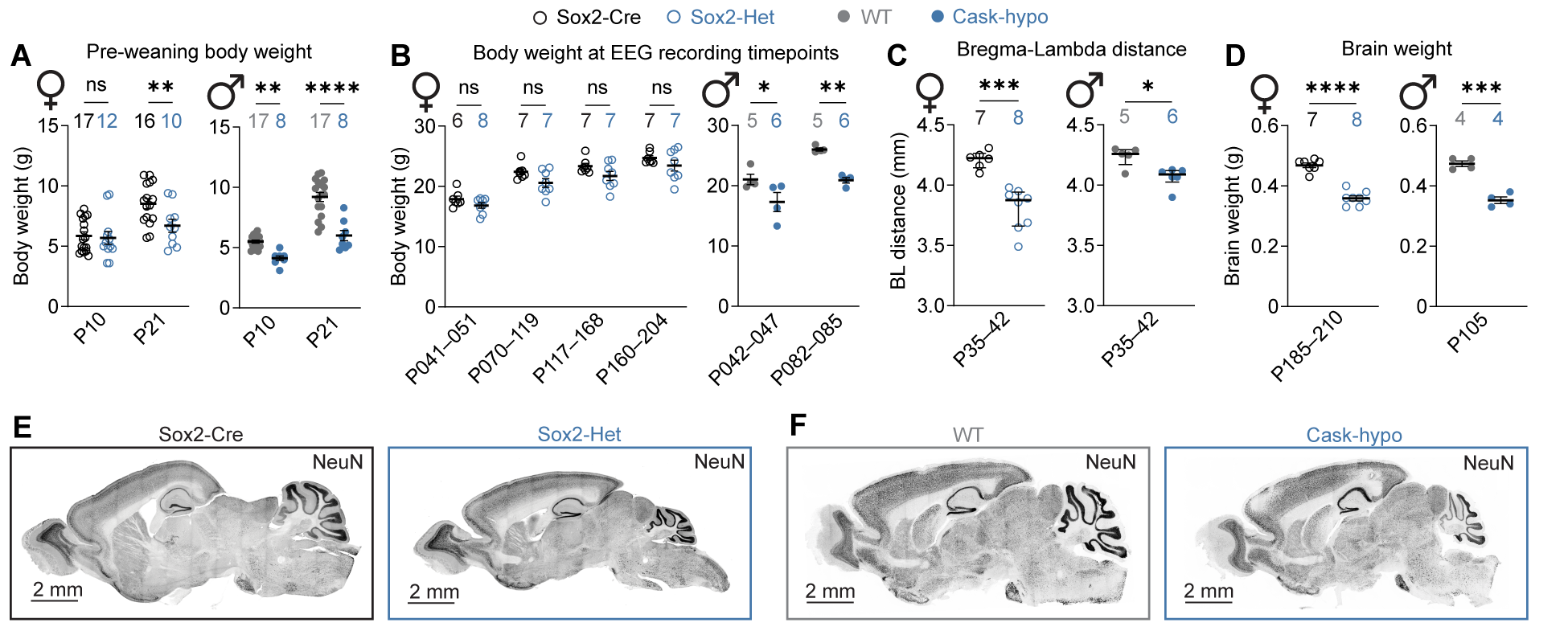

### Supplemental figure 2

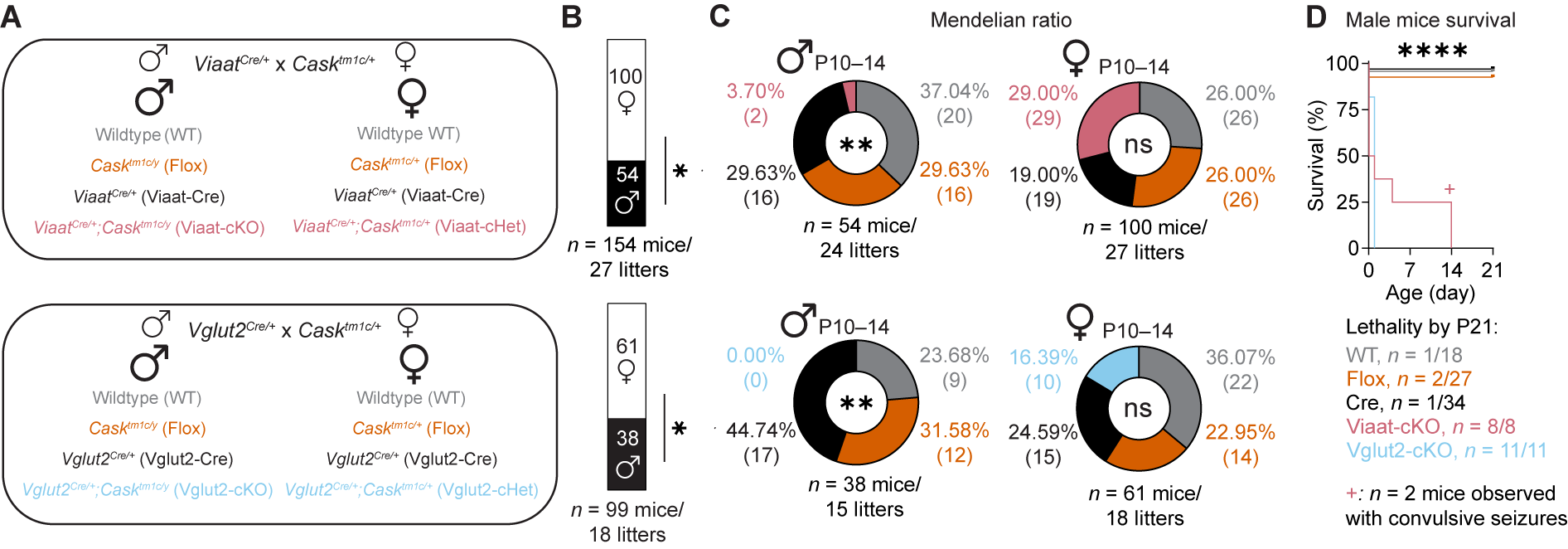

### Supplemental figure 3

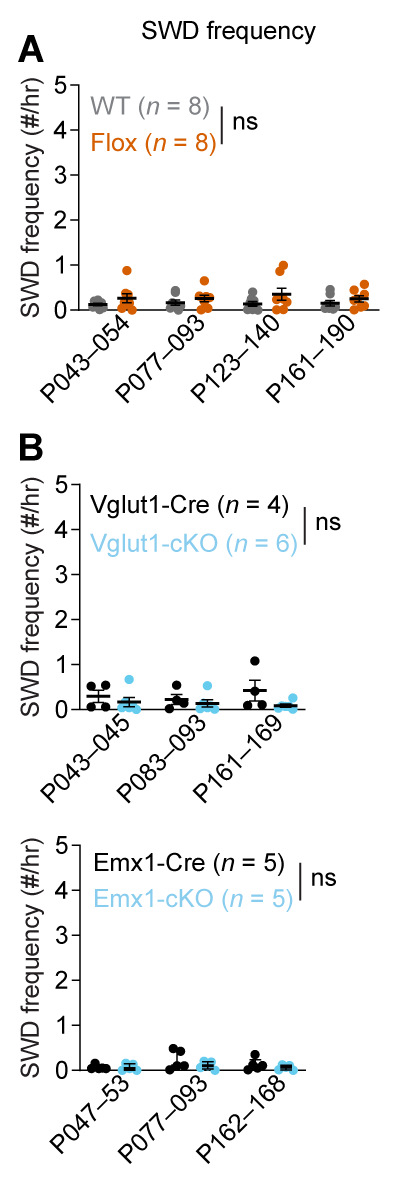

### Supplemental figure 4

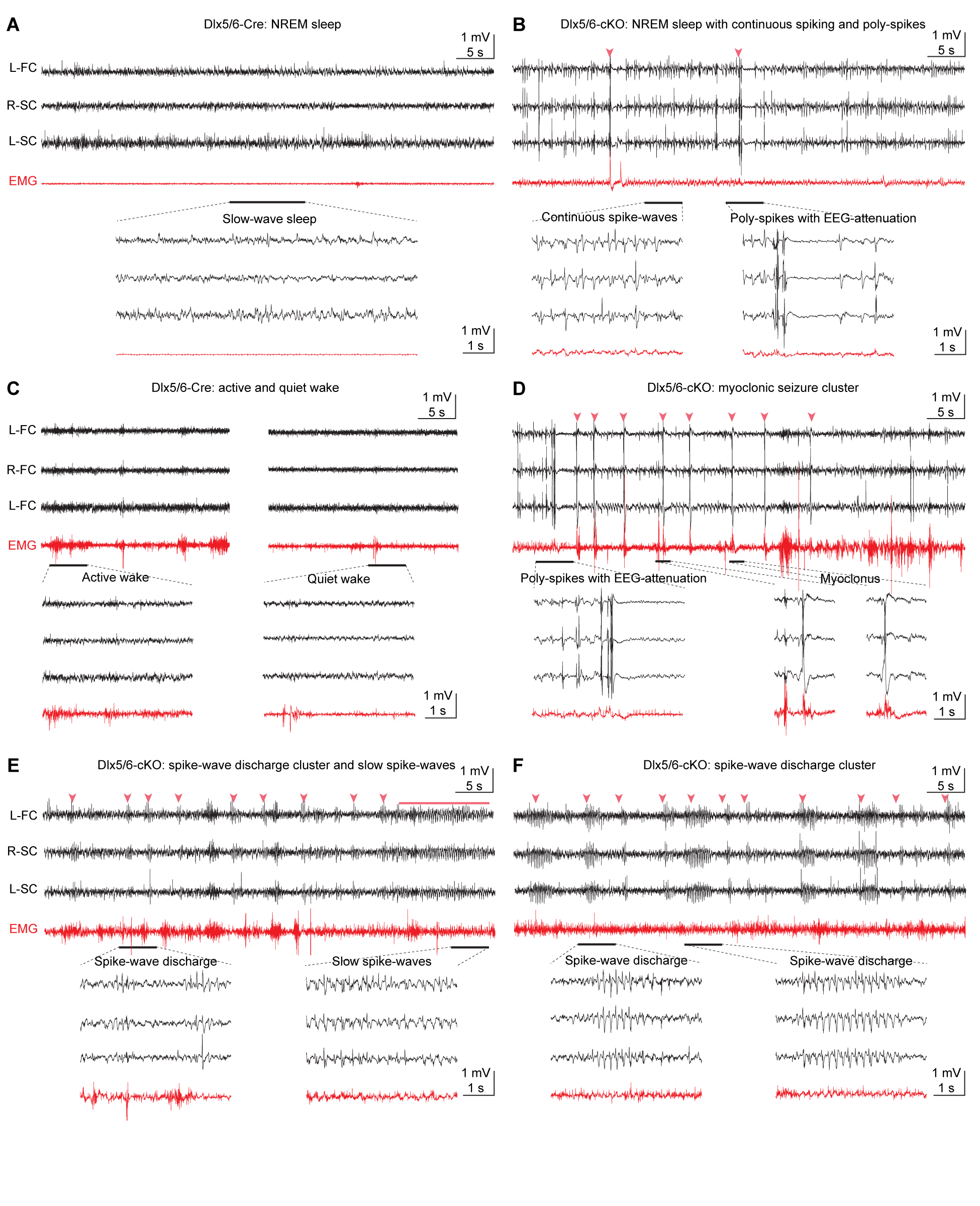

### Supplemental figure 5

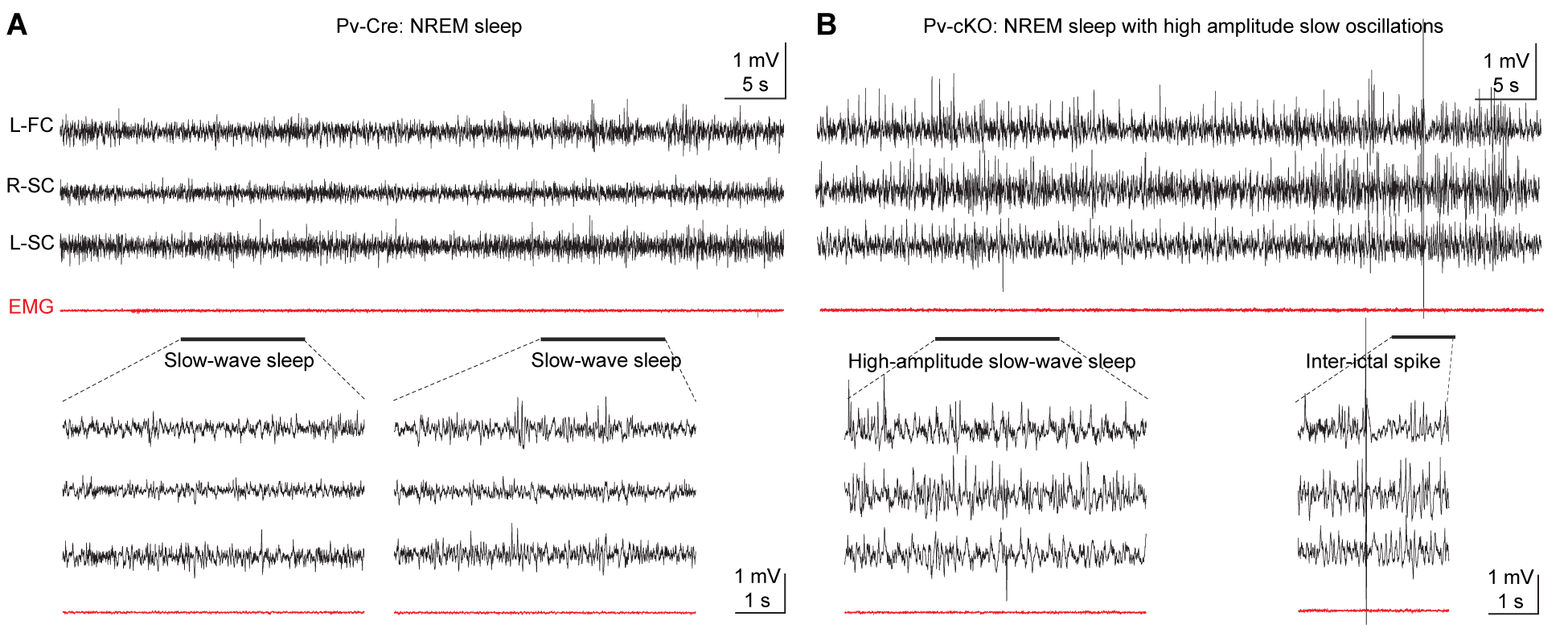

### Supplemental figure 6

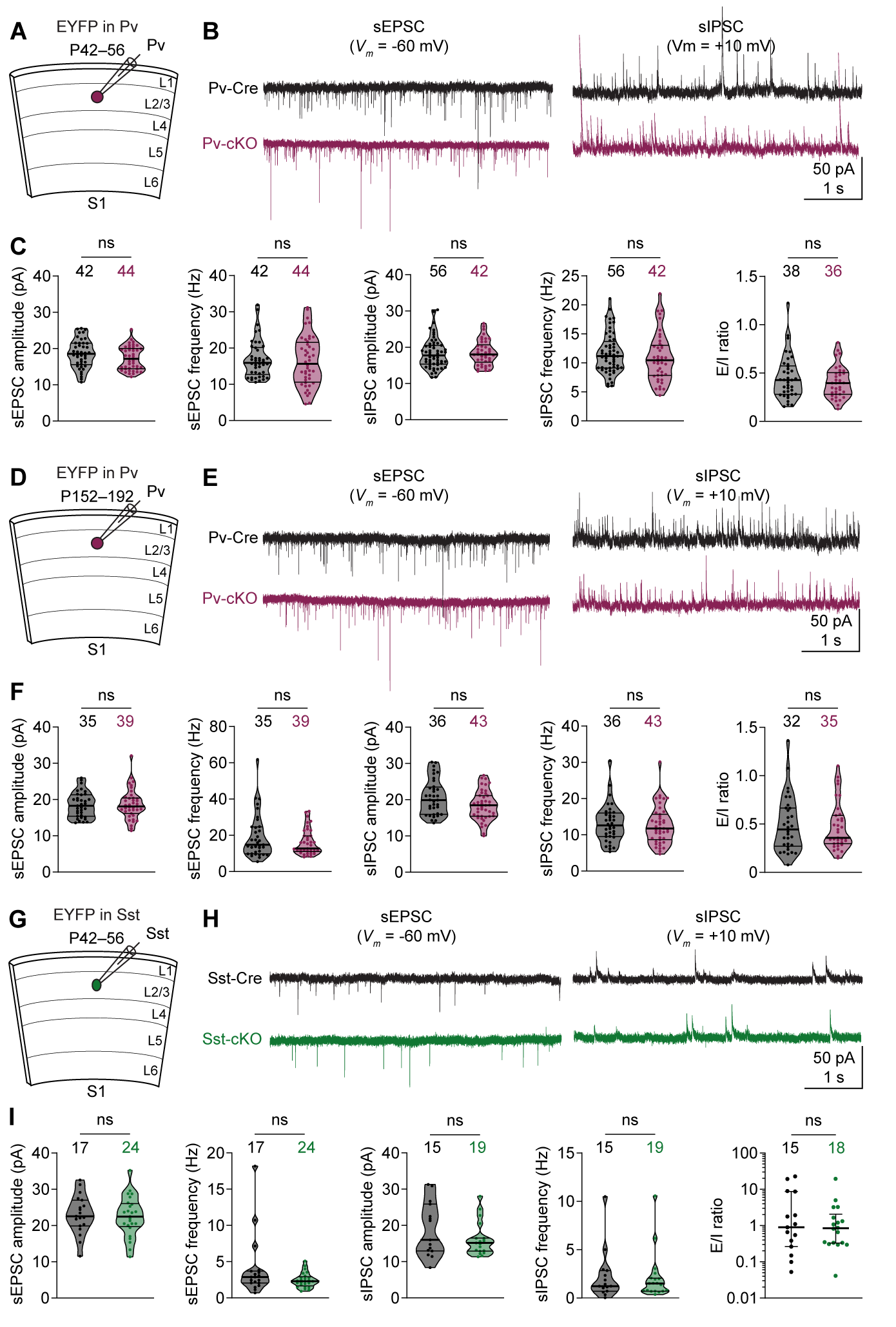
